# RNAtomy: A Structure-Aware RNA Foundation Model for Transferable Structural and Functional Representations

**DOI:** 10.64898/2026.11.10.765792

**Authors:** Azam Shirali, Ishaan Garg, Andrey Vasilyev, Pavitra Soni, Parshatd Govindasamy, Kalai Mathee, Prem Chapagain, Giri Narasimhan

## Abstract

RNA plays a critical and diverse role in gene expression, catalysis, regulation, and molecular recognition. This functional versatility arises from the interplay between nucleotide sequence, structural organization, and cellular context. Nevertheless, most RNA foundation models are pretrained primarily using sequence or secondary-structure information, while the systematic integration of three-dimensional geometric and conformational features remains comparatively underexplored.

Here, we introduce **RNAtomy**, a self-supervised **RNA** foundation model designed to learn the structural ana**tomy** of RNA from inter-nucleotide geometry and residue-level conformational descriptors. Its design has three main components: 1) joint masked reconstruction of pairwise geometry and residue-level descriptors to connect spatial organization with local conformation; 2) a hybrid Mamba–Transformer backbone in which Mamba blocks efficiently propagate global context, while shifted-window attention compensates for the absence of explicit content-dependent token interactions in Mamba by modeling relationships within local regions of the pairwise map; and 3) axis-aware feature-wise linear modulation that maps nucleotide-level conformations to the row and column axes of pairwise geometry.

We assess **RNAtomy** across seven diverse tasks involving structural and functional predictions at the nucleotide, nucleotide-pair, and RNA levels. **RNAtomy** achieved particularly strong performance on structural tasks, outperforming established RNA foundation models and task-specific approaches in contact and long-range base-pair prediction. Its representations also transferred effectively to interaction-related tasks, despite the substantial differences between these applications and the structural pretraining objective. On functional tasks, **RNAtomy** achieved superior performance in translation-related prediction and regulatory-element identification. Notably, **RNAtomy** achieved these results despite being pretrained on a substantially **smaller training set** than the RNA foundation models used as baselines. Together, these results highlight how rich structural information, when learned through an expressive architecture, can support transferable RNA representations across biologically distinct structural and functional applications.

## 1 Introduction

Ribonucleic acid (RNA) is central to gene expression, connecting genetic information encoded in DNA to protein synthesis while also participating directly in catalysis and regulation. Ribosomal and transfer RNAs support translation, small nuclear and nucleolar RNAs contribute to RNA processing and modification, and diverse regulatory RNAs control gene expression [5,4]. Self-splicing RNAs exemplify RNA’s catalytic capacity, whereas riboswitches couple ligand recognition to regulatory responses [19,52]. In messenger RNAs, untranslated regions and interactions with RNA-binding proteins further shape the regulation of protein production [31,39]. The functional versatility of RNA arises from the interplay among its nucleotide sequence, structural organization, and molecular environment. Therefore, computational models that exploit the relationship between sequence and structure are essential for predicting RNA properties across various structural and functional tasks.

The work on RNA foundation models such as RNA-FM, RiNALMo, and AIDO.RNA [7,28,63] suggested that pretraining with larger models and larger data sets can produce embeddings that support diverse RNA tasks. Subsequent models pushed this paradigm farther by enriching it through homologous sequence alignments, motif-aware masking, adaptive tokenization, or relative positional encoding [59,47,44,14]. Although these models appear to learn base-pairing patterns, such structural signals may have emerged from sequence relationships rather than from explicit structural features such as inter-nucleotide distances or relative orientations. Also, their structural generalization remains challenging in low-homology and cross-family settings [58]. To address these limitations, several RNA pretraining approaches have introduced structural information more directly. RNABERT [2] combined masked-language modeling with structural-alignment learning; MP-RNA [55] incorporated predicted secondary structure; and ERNIE-RNA [57] introduced base-pairing priors into its attention mechanism. OmniGenome [54] aligned sequence and secondary-structure representations through complementary sequence-to-structure and structure-to-sequence objectives, while structRFM [62] used predicted canonical base pairs to coordinate the masking of paired nucleotides. These studies high-lighted the importance of structural supervision; however, the methods used mainly represent RNA structure through paired states, dot-bracket annotations, contact patterns, or attention biases. These approaches do not fully capture the distances, orientations, and local backbone conformations associated with RNA’s three-dimensional (3D) structure. But Xu et al. [53] showed that models supplied with 3D representations often outperform sequence-only alternatives, particularly when labeled downstream data are limited.

Unfortunately, leveraging this advantage during pretraining is challenging because experimentally determined RNA 3D structures are significantly underrepresented relative to proteins in structural databases [48,42] and are less diverse compared to RNA sequences. Fortunately, powerful folding methods offer a practical solution to bridge this gap. Tools like trRosettaRNA, RhoFold+, and DRfold2 [48,32,22] can derive 3D structural information from sequences and act as feature generators, transforming large collections of sequences into detailed structural information to guide pretraining models. Motivated by these advances, we extracted structure-derived conformational descriptors for pretraining.

From an architectural perspective, most RNA foundation models use Transformer encoders, whose self-attention mechanism directly models content-dependent relationships but has computational and memory costs that grow quadratically with the number of tokens [46]. Mamba-based architectures reduce the time and space costs from quadratic to linear, while achieving competitive performance. In some cases, Mamba-based solutions such as BiMba [33] and PUMBA [34] outperformed their corresponding Transformer-based solutions [10,38]. To compensate for the lack of an attention mechanism in Mamba, Hatamizadeh et al. [17] successfully integrated attention blocks into a Mamba backbone while retaining all its other advantages. HydraRNA [21] employed that hybrid architecture and showed improvements over pure Transformer RNA language models across multiple sequence-related tasks.

Here, we introduce **RNAtomy**, a self-supervised **RNA** foundation model designed to learn the structural ana**tomy** of RNA from inter-nucleotide geometry and residue-level conformational descriptors. **RNAtomy** is based on a hybrid Mamba–Transformer architecture that learns a shared representation of pairwise geometry maps and residue-level conformational and structural-context descriptors. The key contributions of this work are as follows:

**– New Features:** We formulate RNA pretraining by integrating geometry maps with explicit residue-level descriptors of backbone conformations, glycosidic torsions, and phosphate placement, incorporating new features into RNA representation learning.

- **Self-supervised Pre-training: RNAtomy**’s multimodal framework links residue-level backbone conformations to ordered inter-nucleotide geometry. A key contribution is the joint masked reconstruction of features to learn a unified structural representation during self-supervised pretraining.
- **Hybrid Architecture: RNAtomy** introduces an efficient, vision-inspired structural-map encoder built on a hybrid Mamba–Transformer architecture. The encoder converts pairwise RNA geometry into patch tokens and combines bidirectional Mamba blocks for global context propagation with shifted-window Transformer blocks for modeling relationships within structural neighborhoods.
- **Diverse Downstream Applications: RNAtomy** learns transferable backbone representations through multimodal pretraining. The model extracts valuable structural knowledge to support both **structural and functional** applications for RNA. Superior performance across diverse RNA-related tasks over existing approaches is presented on several benchmarks.
- **Training Set Size: RNAtomy** was pretrained with a substantially smaller dataset, highlighting the power of the architecture.

## 2 Methods

### 2.1 Pretraining Data and Input Representation

We prepared the pretraining dataset from RNAcentral [45]. To reduce redundancy and better represent less common RNA types, we sampled sequences by RNA category and clustered them with CD-HIT-EST [11] at 95% sequence identity, ensuring at least 80% alignment coverage. The resulting candidate collection comprised 391,845 non-redundant sequences across 16 RNA categories. We selected RNAs ranging from 50 to 1024 nucleotides in length for which we successfully generated the structural features and met our quality control standards. Appendix A.1 provides detailed procedures for sequence selection, clustering, and quality control.

Each RNA of length *L* is represented using two complementary structural inputs Fig. 1-A. The first is a pairwise geometry tensor **X** ∈ ℝ^4*×L×L*^, comprising one inter-nucleotide distance-derived channel and three orientation-derived channels generated by trRosettaRNA [48]. These maps were predicted from multiple-sequence-alignment features and a SPOT-RNA secondary-structure estimate [35] and therefore provided spatial information. Long RNAs were processed using overlapping windows, with predictions for shared residue pairs averaged when the windows were assembled into a full-length map. Because relative orientations depend on the ordering of the two nucleotides, we treat the orientation channels as directional rather than symmetric.

**Fig. 1:**
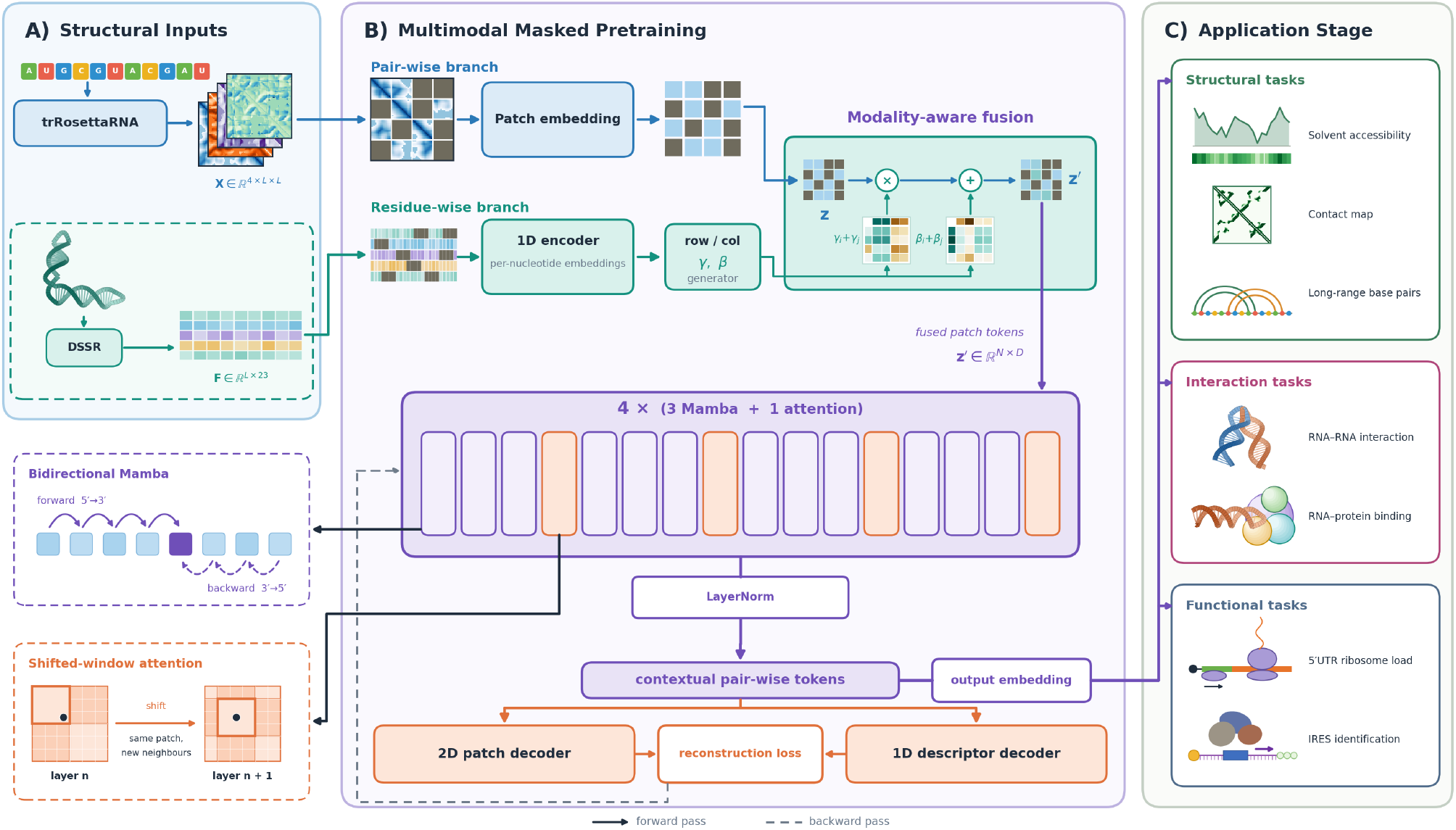
Overview of RNAtomy. **A)** Pairwise geometry maps from trRosettaRNA and residue-wise descriptors from DSSR provide complementary structural inputs. **B)** The two modalities are combined through axis-aware FiLM conditioning and processed by a hybrid bidirectional Mamba–Transformer encoder using joint masked-reconstruction objectives. **C)** The pretrained representations are adapted to structural, interactional, and functional RNA-related tasks.

The second input is a residue-wise descriptor matrix **F** ∈ ℝ^*L×*23^, extracted using DSSR [25] from predicted 3D RNA coordinate models. The 23 features comprised sine–cosine encodings of the six backbone torsions *α, β, γ, δ, ϵ, ζ* and the glycosidic torsion *χ*, two continuous phosphate-geometry descriptors (<monospace>ssZp </monospace>and <monospace>Dp</monospace>), and seven binary indicators describing pairing and membership in stems, helices, junctions, hairpins, bulges, and internal loops (see Appendix A.2 for more details).

The local conformation influences how the RNA backbone positions individual bases for pairing, stacking, ligand recognition, and interactions with other biomolecules. Coordinated combinations of backbone torsion angles recur across RNA structural motifs [30]. The retained backbone angles and glycosidic torsion therefore describe local conformational properties that may not be uniquely recoverable from the selected pairwise geometry maps. The phosphate-placement descriptors provide complementary information related to ribose pucker, while the binary annotations situate each nucleotide within its secondary-structure context [8,25].

These inputs together provide complementary insights into RNA organization: **X** represents spatial relationships between nucleotide pairs, while **F** characterizes the local conformation and structural context of individual nucleotides.

### 2.2 RNAtomy Architecture

**RNAtomy** uses a structure-aware encoder built around a hybrid Mamba–Transformer architecture. An overview of the model is shown in Fig. 1. The standardized pairwise geometry map is divided into non-overlapping *P* × *P* patches and projected into *D*-dimensional tokens by a convolutional patch-embedding layer producing *N* = ⌈*L/P*⌉^2^ pairwise tokens for an RNA of length *L*. Two independent learned positional-embedding tables encode the row and column coordinates of each patch. This axis-specific encoding distin-guishes pairs (*i, j*) and (*j, i*) and is important for directional features, such as distinguishing between forward and reverse orientation of nucleotide *j* in the local coordinate frame of nucleotide *i*, Fig. 1-B.

The tokens are processed by a core encoder that uses a 16-block hybrid backbone arranged as four repeated groups of three bidirectional Mamba blocks followed by one shifted-window Transformer block, thus containing 12 Mamba blocks and four window-attention blocks. Each Mamba block is bidirectional, allowing scans of the token sequence in both directions and for the information to propagate across the complete pairwise representation while reducing the directional bias introduced by flattening a 2D map. The shifted-window attention blocks complement this global propagation by modeling content-dependent relationships within local regions of the pairwise representation and, through alternating window shifts, communicating information across window boundaries [15,24].

A nucleotide-level encoder processes the residue-wise descriptors separately and summarizes them over sequence segments aligned with the pairwise patch grid. **RNAtomy** then uses axis-aware feature-wise linear modulation (FiLM [29]) to incorporate these summaries into pairwise tokens. Independent transformations for the specific columns and rows generate scale and shift parameters, giving the conditioned token at the pairwise position (*i, j*) as

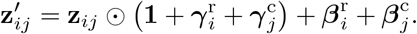

This formulation connects the local conformations of the two participating sequence regions to their pairwise representation while allowing residue-wise information to affect the ordered row and column axes differently. The encoded features can subsequently be retained as a 2D grid for nucleotide-pair tasks, aggregated along one map axis for nucleotide-level tasks, or pooled over valid patches for whole-RNA tasks. Additional architectural details are provided in Appendix B.

### 2.3 Self-Supervised Pretraining

**RNAtomy** is pretrained through joint masked reconstruction of the pairwise and residue-wise structural representations. For pairwise-map reconstruction, we sample a masking ratio uniformly between 0.45 and 0.65, and replace the selected valid patches with a learned mask token. The encoder reconstructs the standardized distance and orientation values of each masked patch. A signal-aware weighting scheme assigns greater weight to patches with stronger geometric signals, reducing the objective’s domination by the comparatively abundant background regions.

For residue-wise reconstruction, we select approximately 15% of valid nucleotide positions in contiguous spans and replace their complete descriptor vectors with a learned sentinel. The pairwise encoder output is aggregated along the column axis and interpolated back to nucleotide resolution to reconstruct the masked descriptors. We use circular loss functions for the sine–cosine torsion representations, a robust regression loss function for the continuous geometric descriptors, and binary cross-entropy for the structural indicators (Appendix B.5). The complete loss function is given by:

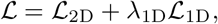

where ℒ_2D_ and ℒ_1D_ denote pairwise and residue-wise reconstruction losses, respectively, and *λ*_1D_ = 0.5.

During pretraining, we withhold the complete residue-wise input for a subset of samples. For these examples, FiLM conditioning reduces to the identity transformation, and the model reconstructs the residue-wise descriptors from the pairwise representation. This modality-dropout strategy improves robustness when residue-wise descriptors are unavailable and encourages the pairwise branch to learn features that support recovering local nucleotide conformations.

Training additionally uses a length curriculum that progressively introduces longer RNAs and shape-aware batching that groups similarly sized pairwise maps while controlling the token budget for long sequences. Appendix B provides detailed masking algorithms, reconstruction losses, batching procedures, optimization settings, and ablation configurations.

## 3 Results and Downstream Tasks

In this section, we first assess the ability of **RNAtomy** to produce biologically informative embeddings by examining how RNAs from different families are organized in its embedding space. We then evaluate the model on seven downstream tasks operating at different levels of resolution, including nucleotide-level regression, nucleotide-pair classification, RNA-pair classification, and whole-RNA prediction. **RNAtomy-FB** is the frozen-backbone transfer setting, in which all parameters of the pretrained encoder remain fixed, and only a lightweight task-specific prediction head is optimized. **RNAtomy** is the full fine-tuning setting, in which the pretrained encoder and prediction head are optimized jointly. This section presents the embedding-space analysis followed by the experimental results for each downstream task. Detailed descriptions of the embedding analysis, prediction heads, optimization settings, loss functions, and additional ablations are provided in Appendix C.

### 3.1 RNA Family Organization in the Embedding Space

To examine the family-level organization captured by RNAtomy, we extracted the embeddings for nine structural RNA families from the ArchiveII benchmark [43] and for families represented in the Rfam 15.0 seed alignments [27]. The encoder produced contextualized patch-token embeddings, which we mean-pooled over valid positions to obtain a 768-dimensional representation for each RNA. We used t-SNE to project these representations into two dimensions and visualize the embedding space, using family annotations only to color the RNA sequences [26]. We quantified family separability in the full space using nearest-neighbor recovery and a cross-validated linear probe. As shown in Fig. 2, **RNAtomy** embeddings organize RNAs according to their family membership. The nine ArchiveII families form largely distinct regions in the embedding space (left). Family-dependent organization is also apparent among the 50 most populated Rfam families and becomes more pronounced for the top 10 families (middle and right). These results indicate that the learned representations retain robust family-level information. Additional visualizations for different numbers of the most populated Rfam families are provided in Appendix C.1.

**Fig. 2:**
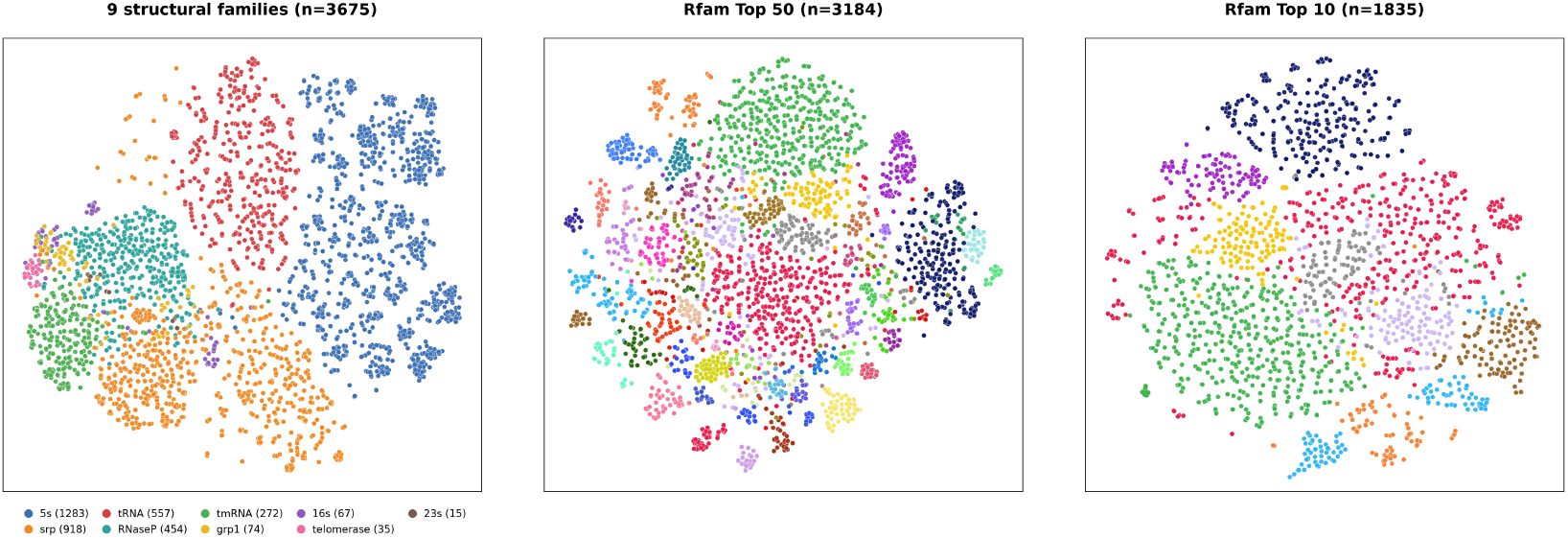
RNA family organization in the RNAtomy embedding space. Two-dimensional t-SNE projections of the embeddings for nine structural RNA families from ArchiveII (left) and the 50 and 10 most populated families in the analyzed Rfam subset (middle and right, respectively). Each point represents one RNA sequence, and colors indicate RNA families.

### 3.2 RNA Structural Properties

#### RNA Solvent Accessibility Prediction

RNA solvent accessibility describes the extent to which each nucleotide is exposed to the surrounding solvent and reflects the influence of its local structural environment. This property is relevant to chemical probing reactivity and interactions with proteins, ligands, and other nucleic acids. As **RNAtomy** is designed to capture transferable RNA structural patterns, we evaluated whether its pretrained representations support nucleotide-level solvent-accessibility prediction using a lightweight regression head. The pretrained backbone was kept fixed and only the prediction head was optimized, corresponding to the **RNAtomy-FB** setting. Appendix C.2 provides details of the prediction head.

We evaluate **RNAtomy** on this task using the RNAsol benchmark’s TR89 training set and TS44 test set [41,56], as provided in the RNAsnap2 release [16]. Each sample contains an RNA sequence with experimentally derived solvent-accessibility values for its constituent nucleotides. We compare against RNAsnap, which uses a support vector machine, and RNAsol, which uses an LSTM together with sequence profiles derived from covariance-model alignments [56,41]. As shown in Fig. 3-A, **RNAtomy-FB** achieved the best performance across all three metrics. Compared with RNAsol, the strongest baseline, **RNAtomy-FB** achieved a PCC of 0.51 versus 0.43, corresponding to an approximately 19% relative improvement. It also achieved an SCC of 0.43 versus 0.42 and a lower MAE of 33.8 versus 34.0 Å^2^. These results indicate that the structural representations learned by **RNAtomy** preserve nucleotide-level information relevant to solvent exposure and support accurate prediction using only a lightweight task-specific head, without fine-tuning the pretrained backbone (See Appendix C.2 for additional results).

**Fig. 3:**
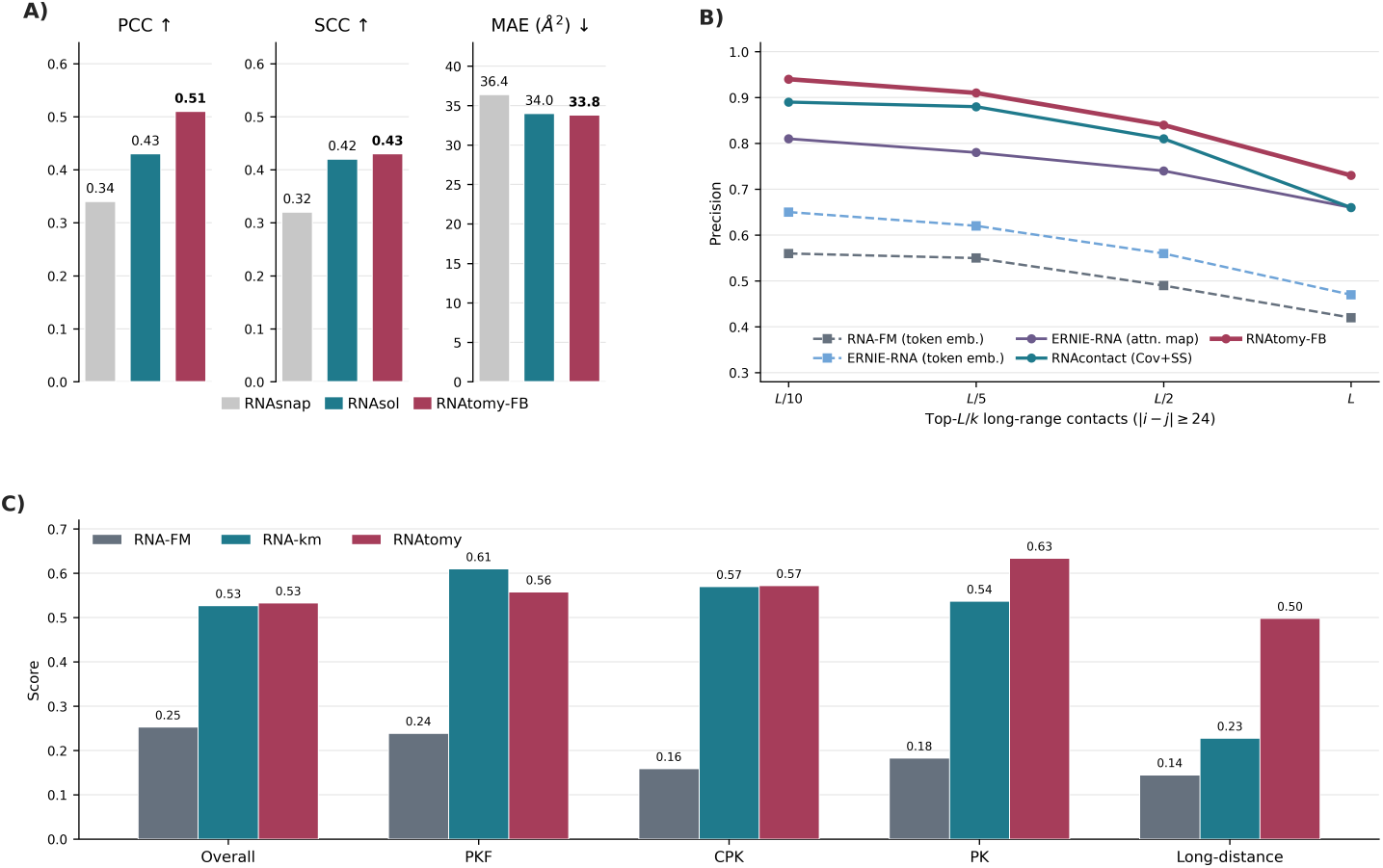
Performance on RNA structural-property tasks. **(A)** Per-nucleotide solvent-accessibility prediction on TS44, evaluated using the Pearson correlation coefficient (PCC), Spearman correlation coefficient (SCC), and mean absolute error (MAE); arrows indicate the preferred direction of each metric. **(B)** RNA 3D contact-map prediction on the RNAcontact benchmark, reported as precision among the top-*L/k* predicted long-range contacts, where |*i™j*| *≥* 24. **(C)** Long-range base-pair prediction on TR0. The bars labeled “Overall” denote the overall F1 score; PKF, CPK, and PK denote prediction accuracy for pseudoknot-free, crossing-pseudoknot, and pseudoknotted base pairs, respectively. Long-distance denotes the reported accuracy for base pairs separated by at least 101 nucleotides.

#### RNA 3D Contact Map Prediction

RNA contact-map prediction determines whether pairs of nucleotides are spatially close in the folded three-dimensional structure. Following RNAcontact [40], two nucleotides are considered to be in contact when the minimum distance between any of their constituent atoms is less than 8 Å. We use the corresponding RNAcontact benchmark, which provides experimentally determined RNA structures for training and testing. The evaluation focuses on long-range contacts with |*i* ™ *j*| ≥ 24, and performance is measured by the precision among the top *L/k* predicted contacts for *k* ∈ {10, 5, 2, 1}. We evaluated the frozen-backbone model **RNAtomy-FB** against RNAcontact, RNA-FM, and ERNIE-RNA [40,7,57]. Appendix C.3 provides details of the prediction head and qualitative comparisons between **RNAtomy**-predicted and ground-truth contact maps.

As shown in Fig. 3-B, **RNAtomy-FB** ranked first at every evaluated cutoff, achieving long-range contact precisions of 0.94, 0.91, 0.84, and 0.73 at *L/*10, *L/*5, *L/*2, and *L*, respectively, and exceeding the strongest baseline by 5, 3, 3, and 7 percentage points. Although **RNAtomy** receives predicted pairwise geometry as input, this task evaluates its ability to recover long-range spatial contacts defined from experimentally determined RNA structures. Moreover, **RNAtomy-FB** achieves this performance using a lightweight prediction head, whereas the comparison methods employ substantially deeper supervised residual networks for contact prediction. These findings indicate that the representations learned by **RNAtomy** effectively integrates pairwise geometric information into representations from which long-range 3D contacts can be readily recovered.

#### Long-Range Base-Pair Prediction

Long-range base-pair prediction evaluates whether a model can identify paired nucleotides that are widely separated in the primary sequence and may contribute to the global RNA fold. We formulate the task as pairwise binary classification and define a base pair as long range when |*i* ™ *j*| ≥ 101, following RNA-km [14]. We use the bpRNA-1m partitions released with SPOT-RNA, comprising the TR0 training, VL0 validation, and TS0 test sets after redundancy removal at 80% sequence identity [9,36]. Reference base pairs are obtained from the supplied dot-bracket structures, including the additional bracket types used to represent pseudoknots.

For direct comparison with RNA-km, we follow its RNA-km-style evaluation on the pseudoknotted RNAs in TR0. We compare **RNAtomy** against the published RNA-FM and RNA-km results [7,14]. Details of the probe, decoding procedure, and evaluation protocol are provided in Appendix C.4. The bars labeled “Overall” denotes the F1 score across all base pairs. PKF measures the recovery of pseudoknot-free pairs that do not cross other pairs; PK measures the pseudoknotted pairs whose removal makes the remaining structure pseudoknot-free; and CPK measures the pairs that cross PK pairs.

As shown in Fig. 3-C, **RNAtomy** achieved an overall F1 of 0.53, slightly exceeding RNA-km at 0.52. It also achieved the highest CPK, PK, and long-distance scores, reaching 0.57, 0.63, and 0.50, respectively. The largest improvement occurred for long-distance pairs, for which **RNAtomy** exceeded RNA-km by 27 percentage points. RNA-km remained stronger for PKF pairs, achieving 0.61 compared with 0.56 for **RNAtomy**. Thus, although the improvement was not uniform across all pairing categories, **RNAtomy**’s strong recovery of pseudoknotted and widely separated pairs indicates that its pairwise representations capture global structural relationships that remain accessible without fine-tuning the pretrained encoder.

### 3.3 RNA Interactions

#### RNA–RNA Interaction Prediction

MicroRNAs are small non-coding RNAs that guide Argonaute proteins to complementary sites in mRNAs, commonly within their 3^*′*^ untranslated regions, thereby promoting translational repression or mRNA degradation [3]. We formulate miRNA–target recognition as RNA-pair binary classification: given a miRNA and a candidate mRNA target-site sequence, the model predicts whether they interact. We use the benchmark evaluated by RNAErnie and distributed as MirTar-RAW/DeepMirTar [51,47]. It contains 13,860 interacting and 13,860 non-interacting pairs. The two RNAs are encoded independently using the shared pretrained backbone. We evaluate both **RNAtomy-FB** and **RNAtomy** for this task (see Appendix C.5 for details).

We compare against established thermodynamic and alignment-based methods, the task-specific Deep-MirTar model, and pretrained RNA models including RNABERT, RNA-MSM, RNA-FM, and RNAErnie [2,59,7,47]. Fig. 4-A compares all evaluated methods across the five metrics. Full fine-tuning improved **RNAtomy** over **RNAtomy-FB**, increasing accuracy from 0.890 to 0.942, recall from 0.940 to 0.961, precision from 0.810 to 0.824, F1 from 0.876 to 0.887, and AUC from 0.915 to 0.957. Both **RNAtomy** configurations outperformed the four conventional methods across every metric. **RNAtomy** also surpassed the task-specific DeepMirTar model in accuracy and recall, although DeepMirTar retained higher precision, F1, and AUC. Fig. 4-B focuses on the pretrained RNA models and the two **RNAtomy** configurations. **RNAtomy** achieved the highest recall among these models, narrowly exceeding RNAErnie at 0.958 and demonstrating strong sensitivity to interacting miRNA–target pairs. Notably, **RNAtomy** was pretrained on RNAs ranging from 50 to 1,024 nucleotides, whereas this benchmark pairs miRNAs of at most 26 nucleotides with target-site fragments of at most 53 nucleotides. Its competitive overall performance and highest recall therefore demonstrate transfer across both sequence-length and task-distribution shifts. This generalization is particularly notable because the comparison language models were pretrained on broad non-coding RNA corpora containing short RNA classes, including miRNAs, whose length distribution is closer to that of this benchmark.

**Fig. 4:**
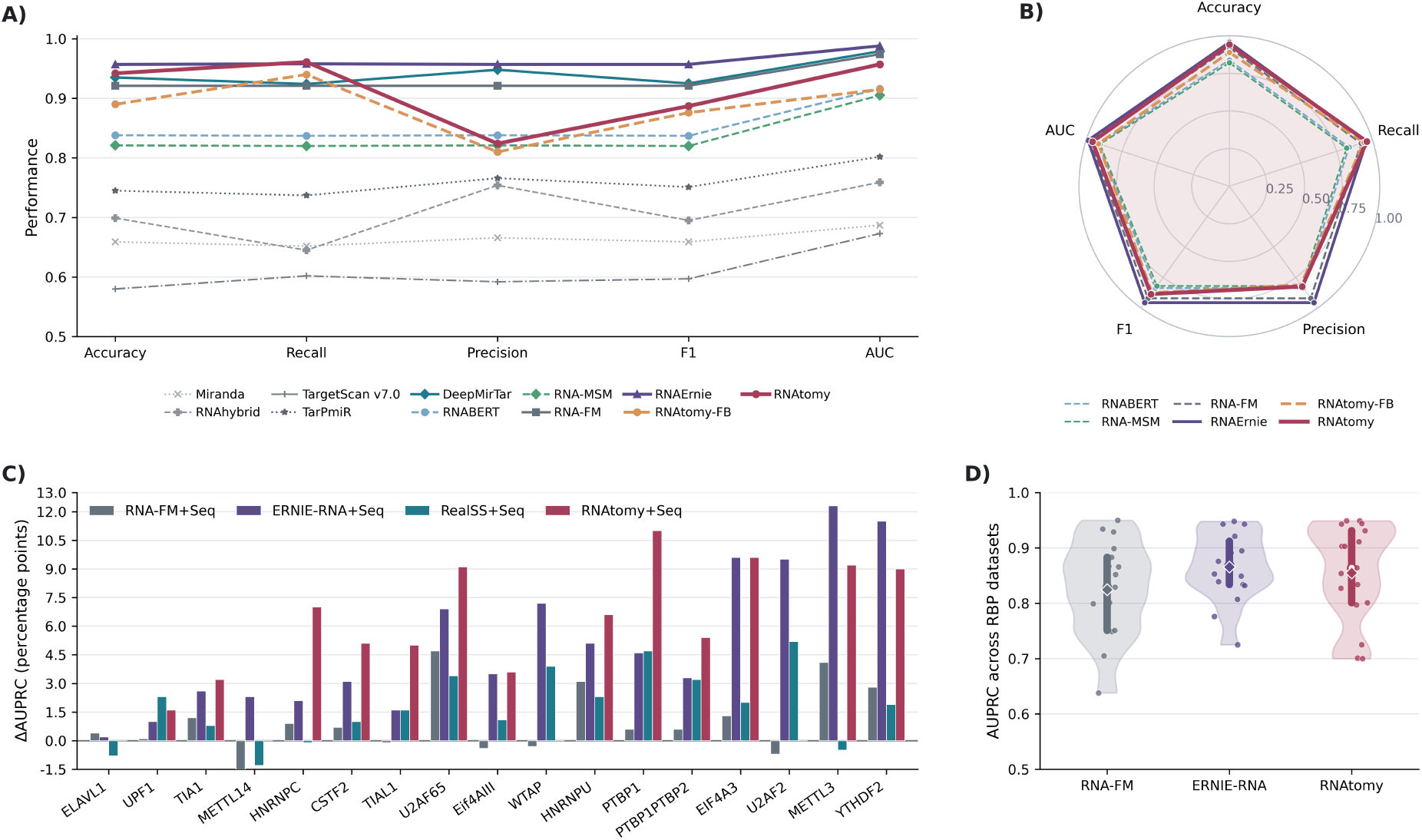
Performance on RNA interaction-related tasks. **(A)** RNA–RNA interaction prediction across accuracy, recall, precision, F1, and AUC. **(B)** Radar comparison of the pretrained RNA models and the two RNAtomy configurations on RNA–RNA interaction prediction. **(C)** RBP–RNA binding performance across 17 RBP-specific datasets, reported as the change in AUPRC, in percentage points, relative to sequence-only PrismNet. **(D)** Distribution of AUPRC values across the 17 RBP datasets for RNA-FM, ERNIE-RNA, and RNAtomy. Individual points represent RBP-specific datasets; thick vertical segments indicate interquartile ranges, open circles indicate medians, and diamonds indicate means.

#### RNA–Protein Binding Prediction

RNA-binding proteins regulate processes including splicing, polyadenylation, localization, stability, and translation. Their recognition of RNA depends on both nucleotide motifs and the structural accessibility of the binding region [13,23,39]. We formulate the benchmark as 17 RBP-specific binary-classification tasks. For each RBP, the model predicts whether a 101-nucleotide RNA window contains a binding site. We use the HeLa-cell datasets from PrismNet, in which each window is associated with a CLIP-seq-derived binding label and an icSHAPE reactivity profile describing its in vivo structural state [37,39]. Following the feature-substitution protocol used by RNA-FM and ERNIE-RNA, the per-nucleotide embeddings from each pretrained model replace the experimental icSHAPE channel and are combined with the one-hot RNA sequence in a matched PrismNet-based classifier [7,57]. The RealSS+Seq configuration is retained as a separate reference that uses the experimentally measured icSHAPE profile. Appendix C.6 provides details of the prediction head and training procedure.

Fig. 4-C shows the change in AUPRC relative to sequence-only PrismNet for each RBP. Incorporating **RNAtomy** representations produced particularly large improvements for PTBP1, EIF4A3, METTL3, U2AF65, and YTHDF2, with gains reaching 11.0 percentage points. These improvements show that the predicted structure-aware embeddings provide binding-relevant information beyond nucleotide identity without requiring experimental icSHAPE measurements as model input. Fig. 4-D summarizes the AUPRC distributions across the 17 RBP-specific datasets. **RNAtomy** achieved a mean AUPRC of 0.855, compared with 0.824 for RNA-FM and 0.865 for ERNIE-RNA. It outperformed RNA-FM on 13 datasets and ERNIE-RNA on 10, while tying ERNIE-RNA on EIF4A3. Although a few larger performance gaps reduced its overall mean below that of ERNIE-RNA, **RNAtomy** performed better or equivalently on most individual RBP datasets. The variation across proteins indicates that the contribution of learned structural information depends on the recognition preferences of each RBP, while the improvements across many datasets support transfer from structure-aware pretraining to RNA–protein binding prediction.

### 3.4 RNA Regulatory and Translational Functions

#### 5^′^UTR Mean Ribosome Load Prediction

Mean ribosome load (MRL) is the average number of ribo-somes associated with an mRNA and provides a quantitative measure of translational activity. The 5^*′*^UTR influences translation through its sequence composition, secondary structure, upstream regulatory elements, and interactions with translation-associated factors. We formulate MRL prediction as whole-UTR regression. We use the variable-length libraries of Sample et al. [31], which contain randomized and natural human 5^*′*^UTRs with experimentally measured MRL values. Random7600 contains 7,600 held-out synthetic sequences from the training distribution, and Human7600 contains 7,600 natural human UTRs and evaluates out-of-distribution generalization.

Because MRL is strongly influenced by sequence-specific elements such as upstream AUGs, the nucleotide-level representations produced by **RNAtomy** are combined with explicit nucleotide identity, positional information, and sequence-derived features describing upstream AUGs, GC content, and UTR length. We evaluate both **RNAtomy-FB** and fully fine-tuned **RNAtomy** using the same task-specific regression head. Details of the prediction head and training procedure are provided in Appendix C.7. We compare against the task-specific Optimus 5-Prime and FramePool models and pretrained RNA models including RNA-FM, UNI-RNA, and ERNIE-RNA [31,18,7,57].

Fig. 5-A shows that **RNAtomy** achieved the highest *R*^2^ on both evaluation sets, reaching 0.95 on Random7600 and 0.88 on Human7600. Fig. 5-B further shows that **RNAtomy** maintained positive performance differences relative to every compared pretrained model on both datasets. Compared with ERNIE-RNA-conv, the strongest pretrained baseline, **RNAtomy** improved *R*^2^ by 3% on Random7600 and 2% on Human7600. As illustrated in Fig. 5-C, full fine-tuning increased performance from 0.89 to 0.95 on Random7600 and from 0.82 to 0.88 on Human7600. **RNAtomy** therefore outperformed ERNIE-RNA-conv despite using fewer parameters, indicating that task-specific adaptation of its structure-aware representation improves both indistribution prediction and generalization to natural human UTRs.

**Fig. 5:**
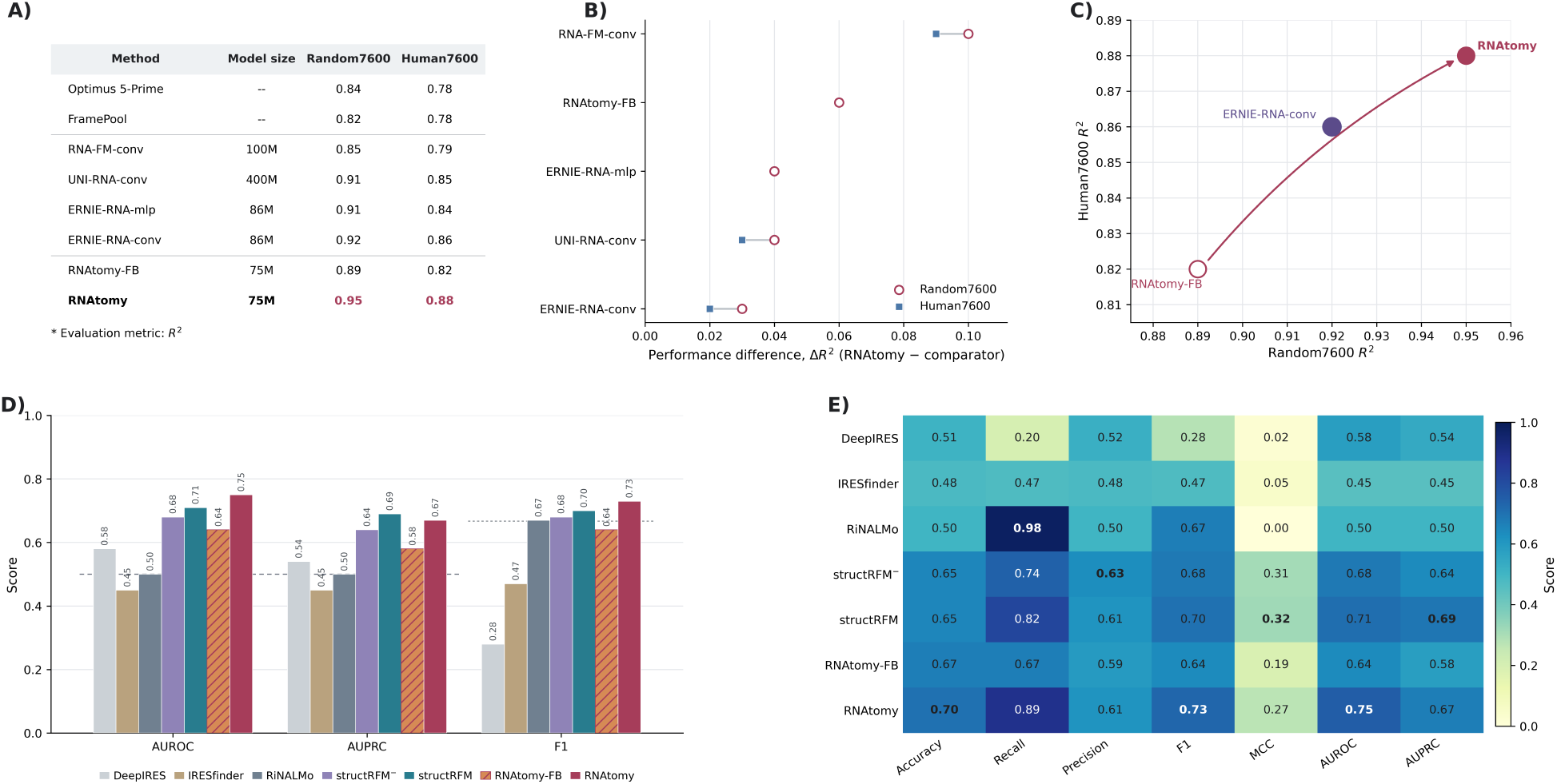
Performance on RNA regulatory and translational tasks. **(A)** MRL prediction performance measured by *R*^2^ on the in-distribution Random7600 and out-of-distribution Human7600 sets. **(B)** Signed performance difference between fully fine-tuned **RNAtomy** and each pretrained comparator, defined as 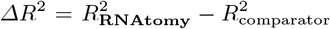. **(C)** MRL performance on Random7600 and Human7600, showing the adaptation from **RNAtomy-FB** to fully finetuned **RNAtomy** and ERNIE-RNA-conv. **(D)** AUROC, AUPRC, and F1 for IRES identification. Dashed reference lines indicate random classification for AUROC and AUPRC (0.5) and the F1 score of an all-positive classifier on the balanced test set (0.667). **(E)** Complete IRES-identification comparison across accuracy, recall, precision, F1, MCC, AUROC, and AUPRC.

#### Internal Ribosome Entry Site (IRES) Identification

IRESs are RNA elements that promote translation initiation independently of the canonical 5^*′*^-cap-dependent mechanism. Their activity depends on both sequence motifs and structural organization. We formulate IRES identification as whole-segment binary classification using the benchmark adopted by structRFM, which was derived from high-throughput reporter assays of human and viral sequences [50,6,62]. Every sample is a 174-nucleotide RNA segment with an IRES or non-IRES label. The nucleotide-level representations produced by **RNAtomy** are combined with the one-hot sequence and supplied to a task-specific classifier. We evaluate both **RNAtomy-FB** and **RNAtomy**. Appendix C.8 provides details of the prediction head, class-imbalance strategy, and training procedure.

We compare against structRFM, its variant without structure-guided pretraining, RiNALMo, and the task-specific IRESfinder and DeepIRES methods [60,61,62]. The baseline values are taken from structRFM. Fig. 5-D shows that **RNAtomy** achieved the highest AUROC, F1, and accuracy, reaching 0.75, 0.73, and 0.70, respectively. Full fine-tuning increased AUROC from 0.64 to 0.75, AUPRC from 0.58 to 0.67, and F1 from 0.64 to 0.73 relative to **RNAtomy-FB**. The complete metric comparison in Fig. 5-E shows that **RNAtomy** also attained a recall of 0.89. Although RiNALMo reached a recall of 0.98, its accuracy and precision of 0.50 and MCC of 0.00 indicate a near-all-positive prediction strategy. **RNAtomy** provided the strongest discrimination by AUROC and the best precision–recall balance by F1, while structRFM retained slightly higher AUPRC and MCC. These results demonstrate that the learned representations transfer effectively to regulatory-element identification and benefit substantially from full task-specific adaptation.

## 4 Discussion

We introduced **RNAtomy** that incorporates key structural features. uses self-supervised pre-training and an improved architecture, resulting in a Foundation Model that performs better on seven different tasks than existing Foundation and task-specific models, even when trained with a much smaller dataset.

While sequence-based models must infer structure implicitly and may miss geometric and conformational relationships that are not readily recoverable from sequence patterns alone, **RNAtomy** learns these structural regularities directly and transfers them across structural, interaction-related, and functional applications. The strongest transfer was observed for structure- and function-related properties, but the representation also supports interaction tasks that differ substantially from masked structural reconstruction.

This structure-to-function transfer reflects how RNA sequence establishes the conformational and spatial contexts required for molecular recognition, regulation, and translation.

**RNAtomy** also transferred across markedly different RNA lengths, from short regulatory regions to RNAs containing thousands of nucleotides, despite **RNAtomy** being trained on a specific range of RNAs. This behavior is supported by its patch-based formulation and its hybrid architecture, which allow the same backbone to process variable-sized pairwise representations while balancing broad context propagation with explicit relational modeling. More generally, the multimodal, modular design enables integration of information at different structural resolutions. Joint learning across pairwise and residue-wise feature levels encourages the model to form a shared structural representation rather than treating each source as an independent auxiliary input. These properties plausibly explain why **RNAtomy** transfers from direct structural properties to biological functions shaped by spatial organization.

Computing structural features is an expensive preprocessing step for the downstream tasks. However, recent progress in computational folding such as AlphaFold3, RhoFold+, trRosettaRNA2, and DRfold2 [1,32,49,22] strengthens the practical utility of **RNAtomy** by serving as scalable feature generators. **RNAtomy** provides a way to reuse this predicted structural knowledge.

Another central finding is that broad transfer does not require relying on scale alone. **RNAtomy** was pretrained on approximately 200,000 RNAs, substantially fewer than the millions of sequences used to pretrain many existing RNA Foundation Models, yet remained effective across structurally and functionally distinct tasks. This result highlights the importance of combining biologically informative inputs with an architecture aligned to their organization. In **RNAtomy**, pairwise geometry exposes spatial relationships, residue-level descriptors capture local conformations, and the hybrid architecture integrates both into transferable representations. Although scale remains valuable, strong structural priors and appropriate architectural design can enable more data-efficient RNA Foundation Models.

An important current limitation is that **RNAtomy** primarily represents each RNA through a static structural description. RNA molecules are intrinsically flexible and may occupy multiple conformational states whose populations change with ions, ligands, molecular partners, and cellular environment. Experimental studies have shown that even well-folded RNAs can explore broad conformational spaces and that changes in ensemble populations can strongly affect binding and cellular activity [20,12]. A single pair-wise map and one set of residue-level descriptors cannot fully represent this heterogeneity. Future work could therefore pretrain on multiple predicted conformers, ensemble-averaged geometric distributions, or condition-dependent structural measurements. Methods that explicitly generate alternative RNA conformations, including trRosettaRNA2, provide a practical starting point for extending **RNAtomy** from static structure representation toward conformational-ensemble learning [49].

This extension would also broaden the model’s application range. Promising directions include predicting mutation-induced structural and functional changes, RNA localization, ligand and protein recognition, RNA– protein complex organization, and the effects of chemical modifications or cellular conditions. The learned representation could additionally be used as a structural prior for RNA sequence design, inverse folding, and generating regulatory or therapeutic RNAs. Such tasks would move beyond recognizing structural patterns toward predicting how RNA structure changes and how it can be intentionally controlled. Overall, **RNAtomy** shows that combining informative structural modalities with a data-efficient and adaptable architecture offers a strong alternative to scale-only pretraining and provides a foundation for connecting RNA geometry to increasingly complex biological functions.

## Code and Data Availability

Source code, pretrained model checkpoints, and pretraining data are publicly available at https://github.com/Azam-Shi/RNAtomy.

## Author contributions statement

G.N. supervised the project. A.S. conceptualized, designed, implemented, and evaluated the model. A.S., I.G., A.V., P.S., and P.G. prepared and preprocessed the datasets. A.S. wrote the first draft. A.S., I.G., and A.V. contributed to writing the appendix. A.S., K.M., P.C., and G.N. reviewed and approved the manuscript. All contributed to interpretations.

## Acknowledgments

Authors thank members of BioRG and Euleris for feedback and comments.

## A Data Preprocessing

### A.1 Pretraining Dataset Preparation

We constructed the pretraining sequence collection from the RNAcentral active-sequence release The RNAcentral Consortium (2021). The source collection contained 40,712,942 sequences. To improve the representation of less abundant RNA classes, sequences were assigned to RNA-type categories by case-insensitive keyword matching of their FASTA descriptions. Sequences without a recognized keyword were assigned to an “other” category. These assignments were used only to guide sampling and were not provided to the model as labels.

Each observed RNA category was initially allocated up to 1,000 sequences, and the remaining sampling budget was distributed in proportion to category abundance. Sampling was performed without replacement using a fixed random seed of 42. To compensate for subsequent redundancy removal, we sampled 640,000 sequences for a target collection of approximately 400,000. The sampled sequences were clustered using CD-HIT-EST at 95% sequence identity, requiring at least 80% alignment coverage for both sequences, and one representative was retained from each cluster Fu et al. (2012).

This procedure produced a candidate collection of 391,845 nonredundant sequences spanning 16 RNA categories. Stratified sampling increased the representation of less abundant RNA types and reduced the proportion of rRNA from 56.47% in the source collection to 38.16% in the candidate collection. The resulting RNA-type distribution is reported in Table S1.

**Table S1:**
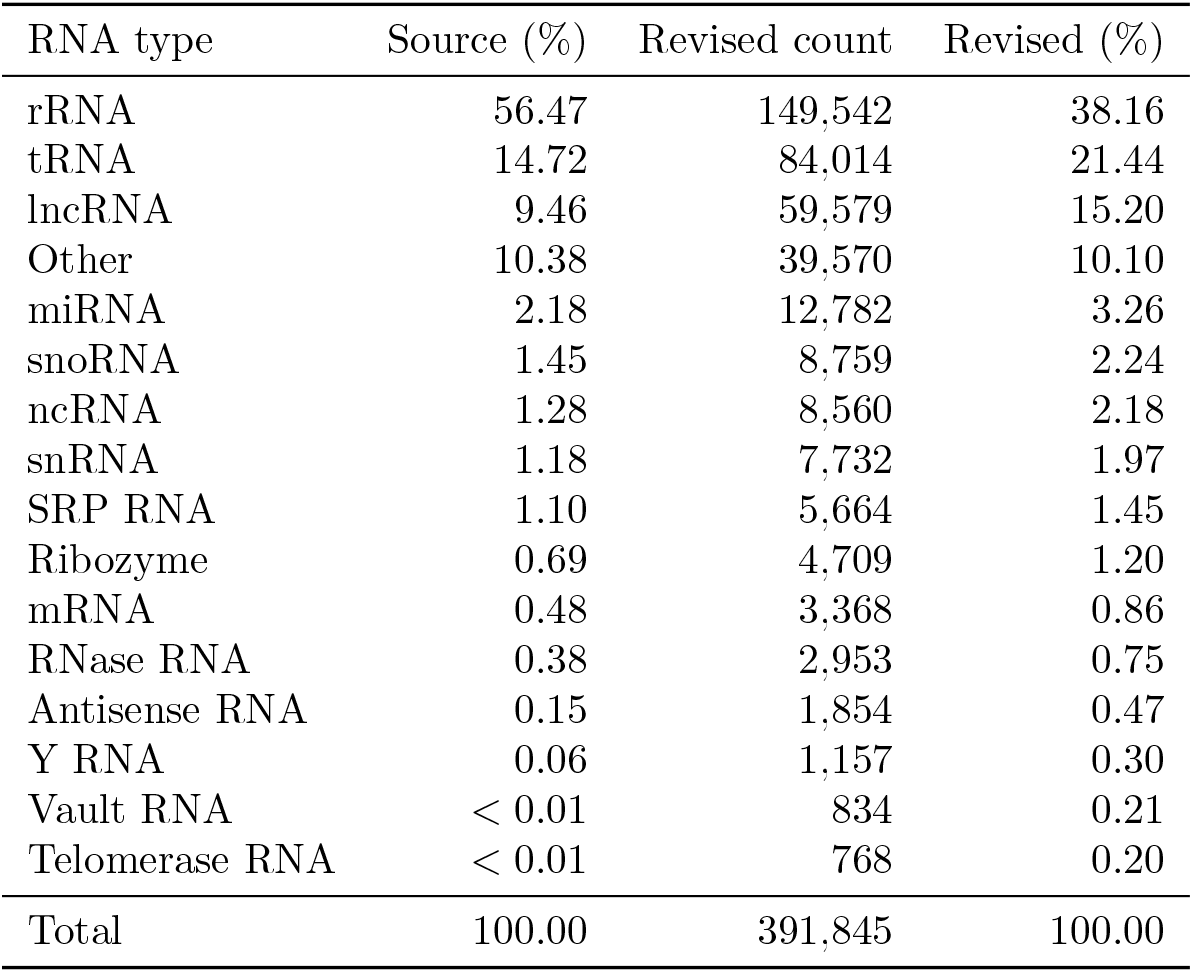
RNA-type distribution in the RNAcentral source collection and the nonredundant candidate collection obtained after stratified sampling and clustering. Categories were assigned by keyword matching of FASTA descriptions and were used only to guide sampling. Percentages may not sum to 100 because of rounding.

| RNA type | Source (%) | Revised count | Revised (%) |
|---|---|---|---|
| rRNA | 56.47 | 149,542 | 38.16 |
| tRNA | 14.72 | 84,014 | 21.44 |
| lncRNA | 9.46 | 59,579 | 15.20 |
| Other | 10.38 | 39,570 | 10.10 |
| miRNA | 2.18 | 12,782 | 3.26 |
| snoRNA | 1.45 | 8,759 | 2.24 |
| ncRNA | 1.28 | 8,560 | 2.18 |
| snRNA | 1.18 | 7,732 | 1.97 |
| SRP RNA | 1.10 | 5,664 | 1.45 |
| Ribozyme | 0.69 | 4,709 | 1.20 |
| mRNA | 0.48 | 3,368 | 0.86 |
| RNase RNA | 0.38 | 2,953 | 0.75 |
| Antisense RNA | 0.15 | 1,854 | 0.47 |
| Y RNA | 0.06 | 1,157 | 0.30 |
| Vault RNA | < 0.01 | 834 | 0.21 |
| Telomerase RNA | < 0.01 | 768 | 0.20 |
| Total | 100.00 | 391,845 | 100.00 |

We subsequently retained sequences between 50 and 1,024 nucleotides in length. The upper limit was determined by the maximum sequence length supported by the structural-feature generation pipeline. Sequences were additionally excluded when pairwise geometry prediction, three-dimensional structure generation, DSSR processing, or other quality-control checks failed to produce the complete set of required input features. After these filtering steps, the final pretraining dataset contained 200,158 RNAs. We randomly assigned 180,142 RNAs to the training set and 20,016 RNAs to the validation set, corresponding to an approximately 90%*/*10% split.

### A.2 Feature Extraction

For an RNA of length *L*, the four outputs selected from trRosettaRNA were organized as a channel-first pairwise geometry tensor:

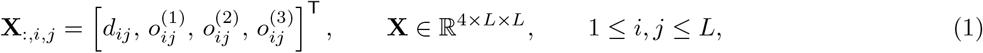

where *d*_*ij*_ is the distance-derived feature and 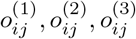 are the three orientation-derived features for nucleotides *i* and *j*. The orientation channels are directional and, in general, 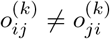.

The residue-wise descriptor matrix **F** ∈ ℝ^*L×*23^ was extracted using DSSR Basic v2.7.0 Lu et al. (2015). The composition and ordering of its 23 features are reported in Table S2.

**Table S2:** Composition and ordering of the DSSR-derived residue-wise features.

| Columns | Descriptor | Encoding |
|---|---|---|
| 1–12 | α, β, γ, δ, ϵ, ζ | Six sine/cosine pairs |
| 13–14 | χ | Sine/cosine pair |
| 15–16 | ssZp, Dp | Standardized continuous values |
| 17 | is_paired | Binary |
| 18–19 | in_stem, in_helix | Binary |
| 20–21 | in_junction, in_hairpin | Binary |
| 22–23 | in_bulge, in_iloop | Binary |

#### Backbone and glycosidic torsion angles

The six backbone torsions describe rotations along the RNA sugar– phosphate backbone, whereas the glycosidic torsion *χ* describes the orientation of the base relative to the ribose sugar. To preserve angular periodicity, each observed angle *a*, reported in degrees, is first mapped to

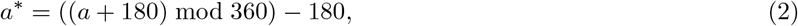

and then represented as

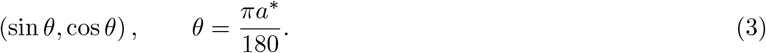

This representation avoids the artificial discontinuity between angles near ™180^*°*^ and 180^*°*^. Undefined angles are represented by (0, 0), which does not correspond to any valid point on the unit circle.

#### Continuous geometric descriptors

The two continuous descriptors characterize the placement of the phosphate group relative to the nucleotide base. For nucleotide *i*, <monospace>ssZp </monospace>is the signed *z* coordinate of the phosphorus atom of the following, 3^*′*^-adjacent nucleotide, expressed in the standard reference frame of the base at position *i*. The <monospace>Dp </monospace>descriptor is the perpendicular distance from this phosphorus atom to the line defined by the glycosidic bond. Both quantities are measured in Å before standardization and provide geometric information correlated with ribose-pucker conformation Lu (a); Chen et al. (2010); Lu (b). DSSR documentation uses approximately 2.9 Å as a practical <monospace>ssZp </monospace>boundary between C3^*′*^-endo-like and C2^*′*^-endo-like conformations in typical cases Lu et al. (2015). We nevertheless retain <monospace>ssZp </monospace>as a continuous feature and do not assign a categorical pucker state from this threshold.

#### Binary structural indicators

The <monospace>is_paired </monospace>indicator is derived from the dot-bracket secondary-structure annotation in the DSSR text output. Chain-separator symbols are removed before indexing, and a nucleotide is labeled as paired when it participates in a matched pair of brackets. The parser recognizes four bracket types, (), [], {}, and ⟨⟩, allowing multiple levels of pairing notation to be represented. Samples are excluded when a readable dot-bracket annotation cannot be recovered or when its nucleotide count does not match the DSSR residue list. The remaining six indicators are derived separately from the DSSR structural objects and denote whether each nucleotide belongs to an annotated stem, helix, junction, hairpin, bulge, or internal loop. These indicators are not mutually exclusive; a nucleotide may therefore receive more than one structural label.

#### Normalization

The sine/cosine encodings are already bounded, and the structural indicators are binary. Consequently, only <monospace>ssZp </monospace>and <monospace>Dp </monospace>are standardized. For each descriptor, the mean and standard deviation are computed once over all observed nucleotide-level values in the training corpus using streaming sums and sums of squares. Each observed value is then transformed using its corresponding training-set statistics, whereas missing values are set to zero after standardization. The angular and binary features are retained on their original scales.

The local conformation influences how the RNA backbone positions individual bases for pairing, stacking, ligand recognition, and interactions with other biomolecules. Coordinated combinations of backbone torsion angles recur across RNA structural motifs Richardson et al. (2008). The retained backbone angles and glycosidic torsion therefore describe local conformational properties that may not be uniquely recoverable from the selected pairwise geometry maps. The phosphate-placement descriptors provide complementary information related to ribose pucker, while the binary annotations situate each nucleotide within its secondary-structure context Chen et al. (2010); Lu et al. (2015). Together, these features complement the pairwise tensor by connecting nucleotide-level conformation with the broader spatial organization learned by the encoder.

## B RNAtomy Architecture and Pretraining

### B.1 Input Representation

Each RNA of length *L* is represented as a 2D tensor

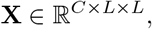

whose *C* channels encode inter-residue geometry (one distance channel and three orientation channels). We additionally provide DSSR-derived Lu et al. (2015) per-nucleotide 1D features

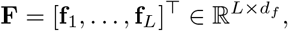

with *d*_*f*_ = 23. Because RNA length varies across samples, within a batch, we zero-pad 2D and 1D features to a common size that is a multiple of the patch size *P*, and a binary validity mask records real versus padded positions; we restrict masking and loss to valid patches throughout.

### B.2 Patch Embedding, Positional Encoding

The input is divided into non-overlapping *P* × *P* patches and projected to dimension *D* by a convolutional patch-embedding layer:

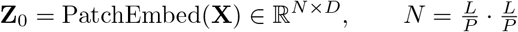

The position of each patch is encoded using two independent learnable tables, **P**^r^ and **P**^c^, for the row and column axes, respectively. A token at patch-grid position (*i, j*) receives

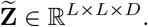

This factorization requires a number of positional parameters that grow linearly with the maximum dimension of the grid and allows (*i, j*) and (*j, i*) to receive different positional representations, which is appropriate for directional orientation channels. During masked pretraining, selected patch embeddings are replaced by a shared learned mask token before positional embeddings and residue-wise conditioning are added. Masked locations therefore retain their pairwise-map coordinates.

### B.3 Hybrid Mamba–Transformer Backbone

The pairwise patch tokens are processed by a 16-block hybrid Mamba–Transformer encoder. Following the MMMT ordering of Liu et al. Liu and Yi (2025), the backbone consists of four repeated groups, each containing three bidirectional Mamba blocks followed by one window-attention Transformer block. The complete encoder therefore contains 12 Mamba blocks and four Transformer blocks.

The Mamba blocks propagate information efficiently across the flattened sequence of pairwise patch tokens. Each block processes the token sequence in both its forward and reverse orders, restores the reverse output to the original ordering, and combines both directions through a pre-normalization residual connection. This bidirectional processing gives every token access to context from both directions and reduces the traversal bias introduced by flattening the two-dimensional pairwise map Gu and Dao (2023); Zhu et al. (2024).

The Transformer blocks use Swin-style shifted-window attention Liu et al. (2021). Attention is restricted to local *M* × *M* regions of the pairwise token grid, providing explicit content-dependent interactions within structural neighborhoods. Alternating non-shifted and half-window-shifted partitions allow information to cross window boundaries. Each block includes two-dimensional relative-position bias, padding-aware attention masks, and a pre-normalization residual MLP. Unlike the hierarchical Swin Transformer, **RNAtomy** does not use patch merging or change the feature resolution between stages; the backbone remains isotropic throughout.

A final layer normalization produces the contextualized pairwise representation

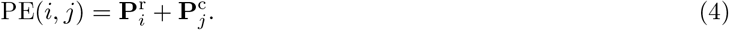

The output can be reshaped to its two-dimensional patch grid for pairwise prediction, aggregated along one map axis for nucleotide-level prediction, or pooled over valid tokens to obtain an RNA-level representation.

### B.4 Residue-Wise Encoding and Axis-Aware Conditioning

The residue-wise descriptor matrix **F** ∈ ℝ^*L×*23^ is first contextualized at nucleotide resolution:

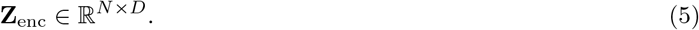

The residue-wise encoder consists of input layer normalization, a linear projection, two residual depthwise-separable one-dimensional convolutional blocks, and a final linear transformation. Padding masks are applied throughout so that padded nucleotide positions do not contribute to the encoded features.

The nucleotide embeddings are averaged over contiguous sequence segments aligned with the pairwise patch grid, producing row- and column-axis summaries:

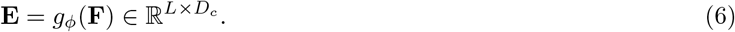

Independent row and column generators transform these summaries into feature-wise scale and shift parameters. The pairwise token at position (*i, j*) is then conditioned using

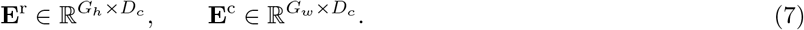

This axis-aware formulation allows the local conformational information associated with the two sequence regions to influence the ordered row and column dimensions independently. Conditioning is applied once, after patch masking and positional encoding, so that the complete hybrid backbone operates on the fused pairwise and residue-wise representation.

### B.2 Joint Multimodal Masked Reconstruction

**RNAtomy** is pretrained by jointly reconstructing masked pairwise geometry patches and masked residue-wise descriptors He et al. (2022); Xie et al. (2022); Bachmann et al. (2022). The complete pretraining objective is

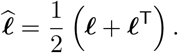

where *ℒ*_2D_ reconstructs the pairwise geometry maps and *ℒ*_1D_ reconstructs the residue-wise descriptors.

#### Pairwise geometry reconstruction

A masking ratio is sampled uniformly from *U*(0.45, 0.65), and the corresponding subset of valid pairwise patches is replaced by a learned mask token. A linear decoder maps each contextualized patch token to the *P*^2^*C* standardized distance and orientation values of its input patch. Reconstruction error is calculated only for masked patches and valid, non-padded pixels.

To reduce domination by weak or nearly uniform geometric regions, masked patches are weighted according to the root-mean-square magnitude of their standardized target values:

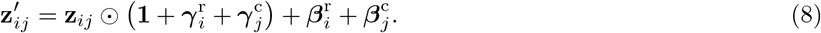

where Ω_*b*_ is the set of masked valid patches for sample *b* and *α*_2D_ = 1. The pairwise reconstruction loss is

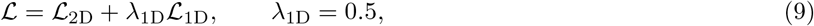

#### Residue-wise reconstruction

Approximately 15% of valid nucleotide positions are selected in contiguous spans of four nucleotides. The complete 23-dimensional descriptor vector at each selected position is replaced by a shared learned sentinel before processing by the residue-wise encoder.

To reconstruct the withheld descriptors, the final pairwise tokens are reshaped to their two-dimensional patch grid and averaged along the column axis using the validity mask. The resulting patch-row representations are interpolated to nucleotide resolution and passed through a reconstruction head that predicts the 23 residue-wise features.

The reconstruction objective follows the three descriptor groups. A cosine loss is used for the seven sine–cosine-encoded torsion angles, Huber loss is used for the two continuous phosphate-geometry descriptors, and binary cross-entropy is used for the seven structural indicators:

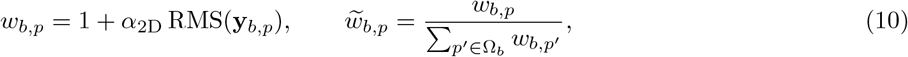

Each component is averaged within its feature group before the three losses are combined. Only valid nucleotide positions and defined descriptor targets contribute to the corresponding loss terms.

#### Modality dropout

To prevent the pairwise branch from becoming dependent on the residue-wise input, the complete residue-wise modality is withheld independently for each sample with probability 0.3. For these samples, the residue-wise embeddings are set to zero, FiLM conditioning reduces to the identity transformation, and a learned no-residue-feature token is added to every pairwise token. The model then reconstructs the residue-wise descriptors from the pairwise representation alone.

For samples retaining the residue-wise input, the reconstruction loss is evaluated at the span-masked positions with *ω*_*bt*_ = 1. For modality-dropped samples, all valid nucleotide positions are reconstructed with *ω*_*bt*_ = 0.5, producing a cross-modal 2D → 1D learning objective. The same no-residue-feature token is used when residue-wise descriptors are unavailable during inference.

### B.6 Pretraining Protocol

Training uses a length curriculum that introduces longer RNAs progressively. During the first 30 epochs, the maximum admitted sequence length increases linearly from the 40th percentile to the maximum of the training-length distribution; all eligible sequence lengths are used thereafter. This curriculum reduces the computational burden during early optimization and allows the model to learn from smaller pairwise maps before processing the largest inputs Bengio et al. (2009).

We use shape-aware token-bucket batching to group RNAs with similar padded map dimensions. Complete groups of equally shaped samples are processed without additional padding, while remaining samples are packed under a token budget. The budget is adjusted for longer RNAs and capped at twice the base budget, balancing GPU utilization against the quadratic growth of the pairwise input size.

#### Optimization and computational resources

Optimization uses AdamW with linear learning-rate warm-up followed by cosine decay. Gradients are accumulated over two mini-batches and clipped to a global norm of 1.0. Exponential-moving-average weights are used for validation and checkpoint selection. **RNAtomy** was pretrained for 150 epochs on a system equipped with four NVIDIA A100 80 GB PCIe GPUs. Pretraining required approximately 85 hours (∼ 3.5 days). The complete hyperparameter configuration is summarized in Table S3.

### B.7 Ablation Study

We evaluate the contribution of residue-wise structural conditioning and model capacity using a 2 × 2 ablation design. Pairwise-only and multimodal configurations are compared at embedding dimensions of 512 and 768. The pairwise-only models use only the inter-nucleotide geometry maps, whereas the multimodal models additionally incorporate the residue-wise descriptors through FiLM conditioning. All other training settings, data splits, and evaluation procedures are held fixed.

**Table S3:** Hyperparameters used for RNAtomy pretraining.

| Hyperparameter | Value |
|---|---|
| Pairwise input channels | 4 |
| Residue-wise input dimension | 23 |
| Patch size | 8 × 8 |
| Embedding dimension | 768 |
| Encoder depth | 16 blocks |
| Backbone arrangement | 4 × (3 Mamba + 1 attention) |
| Bidirectional Mamba blocks | 12 |
| Shifted-window attention blocks | 4 |
| Attention heads | 12 |
| Attention-window size | 16 × 16 patches |
| Pairwise masking ratio | $\mathcal{U}(0.45, 0.65)$ |
| Pairwise weighting coefficient | α<sub>2D</sub> = 1.0 |
| Residue-wise masking ratio | 0.15 |
| Residue-wise mask span | 4 nucleotides |
| Modality-dropout probability | 0.30 |
| Residue-wise loss weight | λ<sub>1D</sub> = 0.5 |
| Mini-batch size per GPU | 16 |
| Gradient-accumulation factor | 2 |
| Effective global batch size | 128 |
| Optimizer | AdamW |
| Peak learning rate | 2 × 10<sup>−4</sup> |
| Weight decay | 0.05 |
| Learning-rate schedule | Linear warm-up and cosine decay |
| Warm-up duration | 20 epochs |
| Training duration | 150 epochs |
| Early-stopping patience | 20 epochs |
| Drop-path rate | 0.1 |
| Gradient clipping | Global norm of 1.0 |
| Training precision | FP32 |
| Length curriculum | 40th to 100th percentile |
| Curriculum duration | 30 epochs |
| Checkpoint evaluation | EMA weights |

**Table S4:** Ablation of residue-wise structural conditioning and embedding dimension on a homology-filtered held-out set of 1,323 RNAs. Performance is measured by pairwise-map reconstruction MSE; lower values indicate better reconstruction.

| Configuration | *d* | Params. (M) | MSE ↓ |
|---|---|---|---|
| Pairwise-only | 768 | 74.4 | 0.18 |
| Pairwise-only | 512 | 33.1 | 0.25 |
| Pairwise + residue-wise conditioning | 768 | 75.6 | **0.11** |
| Pairwise + residue-wise conditioning | 512 | 34.3 | 0.17 |

Residue-wise conditioning consistently improved reconstruction at both model sizes, reducing MSE from 0.18 to 0.11 at *d* = 768 and from 0.25 to 0.17 at *d* = 512. Increasing the embedding dimension also improved both configurations. The best performance was obtained by the 768-dimensional multimodal model, supporting the complementary contribution of residue-wise conformational information and increased representation capacity.

## C Applications

### C.1 RNA Family Organization in the Embedding Space

We evaluated the organization of the **RNAtomy** embedding space using two family-level datasets. The first contains nine structural RNA families from the family-fold partitions of ArchiveII released by Szikszai et al. Szikszai et al. (2022): 16S, 23S, and 5S rRNAs, RNase P, group I introns, signal-recognition-particle RNA, tRNA, telomerase RNA, and tmRNA. Of the 3,865 sequences in this collection, 3,675 had all inputs required by **RNAtomy** and were retained. The second dataset contains the 5,000 longest sequences from the Rfam 15.0 seed alignments Ontiveros-Palacios et al. (2025). Valid **RNAtomy** embeddings were obtained for 4,230 sequences representing 233 of the original 242 families. For the top-*k* analyses, families were ranked by their number of valid sequences in this analyzed Rfam subset.

RNA-level embeddings were extracted directly from the pretrained **RNAtomy** encoder. For each RNA, the contextualized patch-token embeddings were mean-pooled over valid positions, yielding a 768-dimensional vector. RNAs exceeding the encoder’s positional capacity were processed in overlapping windows of 1,024 nucleotides, and the resulting window-level embeddings were averaged to obtain a single representation.

We used t-SNE only to project the 768-dimensional embeddings into two dimensions for visualization van der Maaten and Hinton (2008). The projection was computed using cosine distance, PCA initialization, a perplexity of 60, an early-exaggeration factor of 16, 1,500 iterations, and a random seed of 42. Family labels were used only to color the projected points.

Family separability was quantified in the full 768-dimensional whitened embedding space. Nearest-neighbor accuracy was calculated using leave-one-out 5-nearest-neighbor classification with cosine distance and majority voting. Families containing fewer than five sequences were excluded from the linear-probe evaluation.

Figure S1 shows the embedding-space projections for the 100, 80, 60, 40, 20, and 10 most populated Rfam families. The progressive separation observed as less populated families are excluded is consistent with the quantitative evaluation, indicating that **RNAtomy** retains robust family-level information across diverse RNA classes.

**Figure S1:**
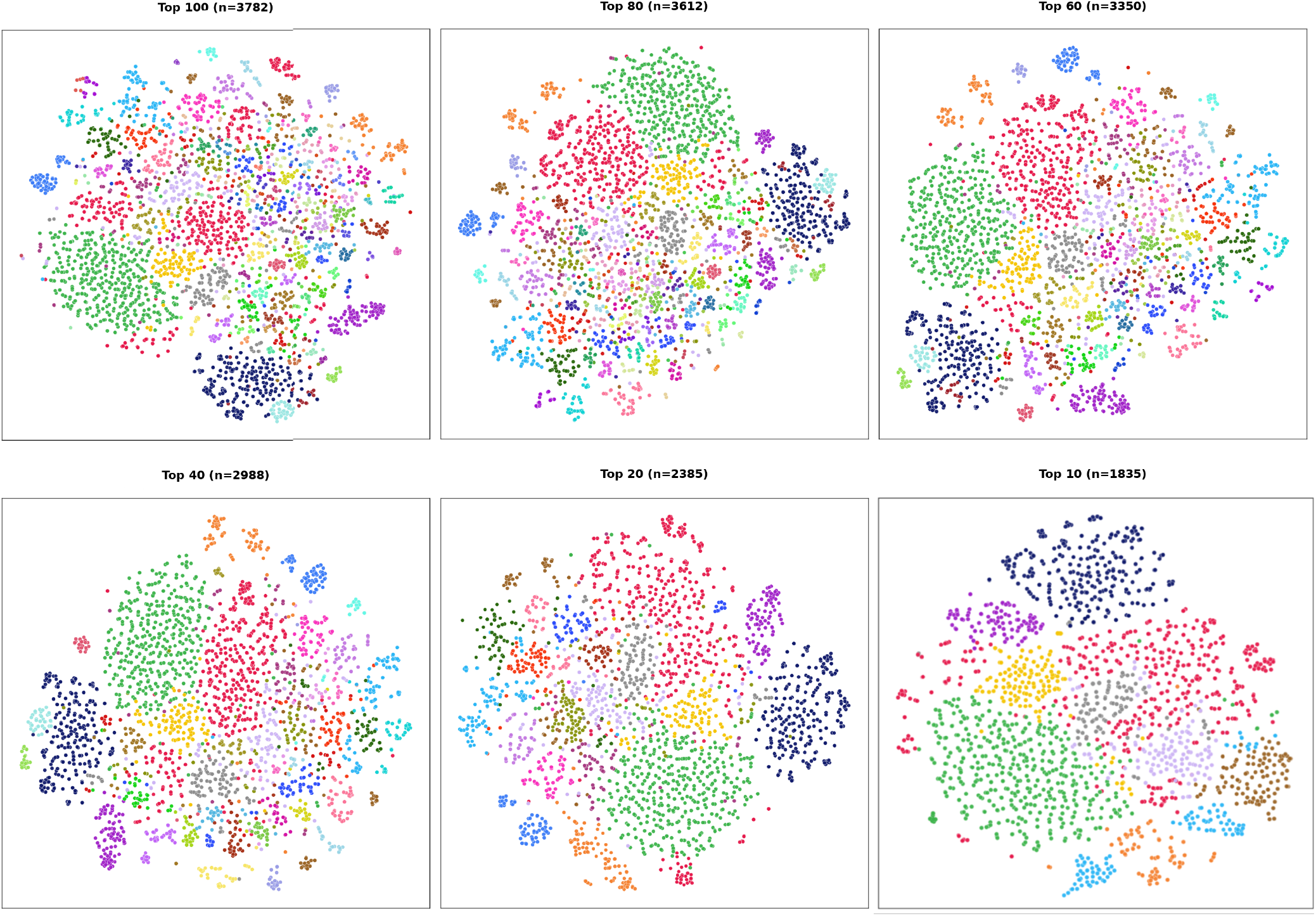
RNAtomy embedding-space organization across Rfam families. Two-dimensional t-SNE projections of the 768-dimensional RNA-level embeddings for the 100, 80, 60, 40, 20, and 10 most populated families in the analyzed Rfam 15.0 subset. Each point represents one RNA, and colors distinguish families within each panel. The projections are used only for visualization; all family-recovery metrics are calculated in the full 768-dimensional embedding space.

### C.2 RNA Solvent Accessibility Prediction

#### Task definition

RNA solvent accessibility describes the degree to which each nucleotide is exposed to the surrounding solvent. This property depends on the nucleotide’s local structural environment, including whether it is located in a paired or unpaired region, a loop, a junction, or another structural motif. Solvent accessibility can influence chemical probing measurements and interactions between RNA molecules and proteins, ligands, or other nucleic acids. We formulate the task as nucleotide-level regression, where the model predicts an accessibility value for every nucleotide in the RNA sequence.

#### Dataset

We use the TR89 training set and TS44 test set distributed with RNAsnap2 Hanumanthappa et al. (2020). These sets were originally constructed for RNAsnap Yang et al. (2017) and were subsequently used by RNAsol for direct comparison Sun et al. (2019). TR89 contains 89 nonredundant protein-bound RNA chains derived from structures with resolution better than 3.0 Å. TS44 contains 44 independent protein-bound RNA chains obtained from structures with resolutions between 3.0 and 4.0 Å. Each RNA is associated with experimentally derived per-nucleotide solvent-accessibility values. We train and select the task-specific prediction head using TR89 and report final performance only on TS44.

#### Task-specific prediction head training

Because solvent accessibility is a nucleotide-level property, a pooled RNA-level embedding does not retain the positional information required for this task. We therefore use the patch-token grid produced by the pretrained encoder and convert it into nucleotide-level representations. For each patch row *g*, we compute an interaction summary by averaging over its valid column patches and retain the corresponding diagonal token:

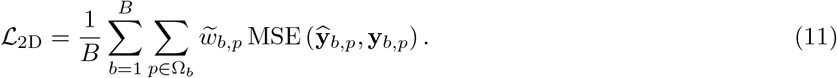

where *C*_*g*_ denotes the set of valid column patches. The averaged representation **r**_*g*_ summarizes the interactions of nucleotide block *g* with the rest of the RNA, while **d**_*g*_ retains its contextualized within-block information. The two representations are concatenated as:

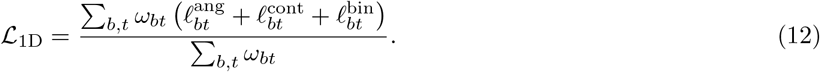

For each RNA *c*, these patch-level representations are linearly interpolated from its valid patch rows to its nucleotide length:

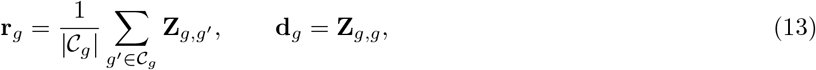

For nucleotide *i*, the interpolated embedding is concatenated with its residue-wise structural descriptor vector:

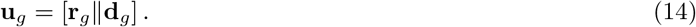

The resulting sequence is processed by a shared lightweight regression head consisting of layer normalization, residual local refinement, and a multilayer perceptron:

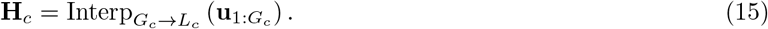

The pretrained backbone remains fully frozen, and only the parameters of *h*_*ψ*_ are optimized. Training minimizes mean-squared error over valid supervised nucleotide positions:

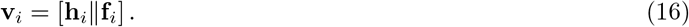

where *m*_*i*_ ∈ {0, 1} is a supervision mask that equals 1 for valid nucleotide positions assigned to the current training window and 0 for padded positions or overlapping positions assigned to another window.

Table S5 reports performance on the TS44 test set using the Pearson correlation coefficient (PCC), Spearman correlation coefficient (SCC), and mean absolute error (MAE).

### C.3 RNA 3D Contact Map Prediction

#### Task definition

RNA contact prediction determines whether two nucleotides occupy nearby positions in the folded three-dimensional structure. The task is formulated as pairwise classification, where each nucleotide pair (*i, j*) is assigned a binary contact label. Long-range contacts connect nucleotides that are distant in the primary sequence and provide information about the global organization of an RNA molecule.

**Table S5:** Comparison with RNAsol, RNAsnap, and pretrained RNA language models on the TS44 test set. The language models use a frozen backbone with a lightweight task head.

| Method | PCC ↑ | SCC ↑ | MAE (Å<sup>2</sup>) ↓ |
|---|---|---|---|
| RNAsol | 0.43 | 0.42 | 34.0 |
| RNAsnap | 0.34 | 0.32 | 36.4 |
| ERNIE-RNA | 0.28 | 0.27 | 49.6 |
| RNAErnie | 0.29 | 0.30 | 43.0 |
| structRFM | 0.31 | 0.30 | 41.7 |
| **RNAtomy-FB** | **0.51** | **0.43** | **33.8** |

#### Dataset

We use the benchmark released with RNAcontact Sun et al. (2021b), which comprises 221 training RNAs (TR221) and 80 test RNAs (TE80) with experimentally determined structures. The downloaded archive contains 211 training structures and the 80 test structures, and we use the structures it contains. Following RNAcontact, long-range contacts are pairs with |*i* ™ *j*| ≥ 24.

#### Task-specific prediction-head training

Contact prediction is an inherently pairwise task, with each target indicating whether nucleotides *i* and *j* are in spatial contact. Following the RNAcontact definition, a pair is labeled as a contact when the distance between their C1^*′*^ atoms is less than 8 Å. Because the pretrained encoder directly produces pairwise representations, its patch-token output **Z** ∈ ℝ^*G×G×D*^ is bilinearly interpolated to nucleotide resolution:

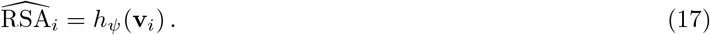

The pretrained encoder is kept frozen during task-specific training (**RNAtomy-FB**), and only a lightweight two-dimensional convolutional prediction head *h*_*ψ*_ is optimized. The predicted contact logits are

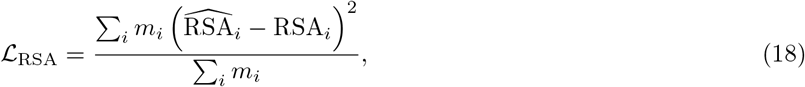

The prediction head applies a channel projection followed by local two-dimensional convolutional refinement with channel-wise layer normalization and GELU activations. Channel-wise normalization is performed independently at each nucleotide pair, while the convolutional operations allow the head to capture neighboring contact-map patterns, including the band-like patterns produced by RNA helices. Symmetrizing the final logits enforces the physical equivalence of contacts (*i, j*) and (*j, i*).

RNA contact maps are highly imbalanced because non-contacting nucleotide pairs substantially outnumber contacts. We therefore train the prediction head using sigmoid focal loss Lin et al. (2017). For a pair (*i, j*), let *p*_*ij*_ = *σ*(*S*_*ij*_), and define

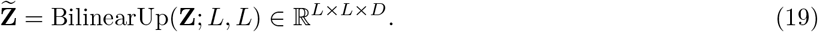

The contact-prediction loss is then

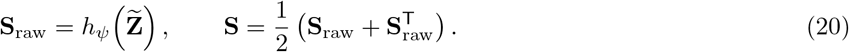

where

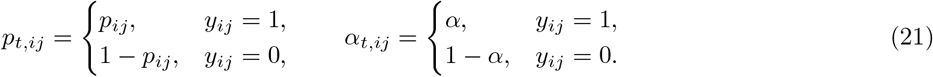

contains the valid supervised pairs in the strict upper triangle, and *m*_*ij*_ excludes invalid or padded positions. We use *α* = 0.5, *γ* = 2, and *s*_min_ = 6. The focal term reduces the contribution of easily classified pairs and concentrates learning on difficult contacts. Restricting supervision to *i < j* avoids counting each symmetric nucleotide pair twice.

For evaluation, long-range contacts are defined by |*i* ™ *j*| ≥ 24, and performance is measured using Top-*L/k* precision for *k* ∈ {10, 5, 2, 1}.

To qualitatively assess contact recovery across different RNA lengths, we visualized ten test RNAs selected by stratified random sampling over the length distribution, without considering prediction performance. As shown in Fig. S2, **RNAtomy**’s predicted contact-probability maps recover the principal long-range contact patterns observed in the experimental structures across diverse RNAs. The visible signal near the diagonal corresponds to short-range nucleotide pairs with |*i* ™ *j*| *<* 24, which are excluded from both labeling and evaluation.

### C.2 Long-Range Base-Pair Prediction

**Figure S2:**
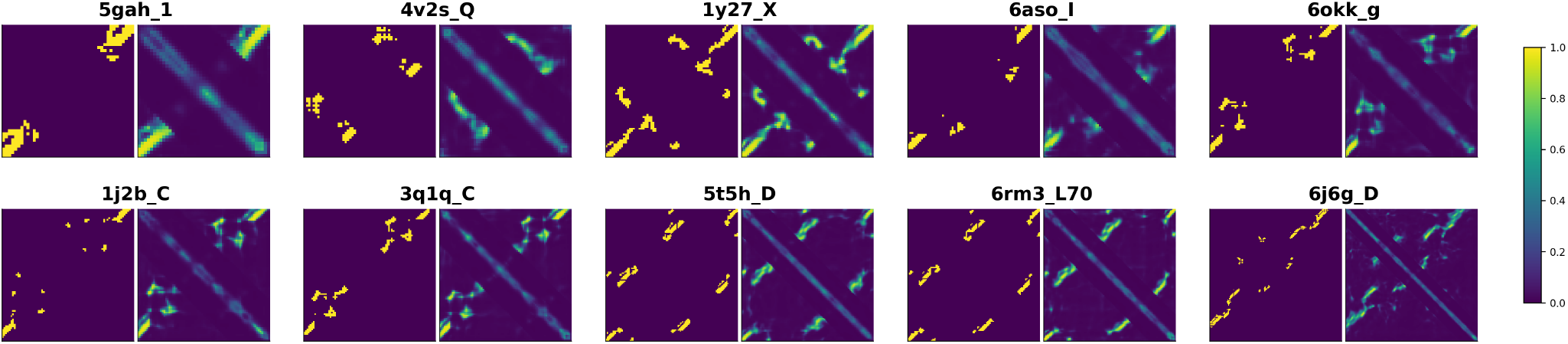
Long-range contact-map predictions for selected samples from the RNAcontact test set. For each RNA, the ground-truth map (left) is shown next to the **RNAtomy** prediction (right). The ground truth is binary (0 or 1) and the prediction is the contact probability, both using the color scale at the right.

#### Task definition

Long-range base-pair prediction evaluates whether a model can identify base pairs between nucleotides that are widely separated in the primary sequence. These interactions are difficult to infer from local sequence context but can strongly influence the global RNA fold. A base pair is classified as long-range when the sequence separation between its two nucleotides is at least 101 positions. We formulate the task as pairwise binary classification over nucleotide positions.

#### Dataset

We use bpRNA-1m Danaee et al. (2018) as partitioned by SPOT-RNA Singh et al. (2019) into training (TR0), validation (VL0), and test (TS0) sets after redundancy removal at 80% sequence identity. TR0 contains 10,814 RNAs, of which 999 contain at least one pseudoknot. Following RNA-km Gong and Bu (2024), a base pair is considered long-range when its two nucleotides are separated by at least 101 positions, |*i* ™ *j*| ≥ 101.

#### Task-specific prediction-head training

The prediction head operates on the encoder’s native two-dimensional representation. The patch-token grid is bilinearly interpolated to nucleotide-pair resolution, producing

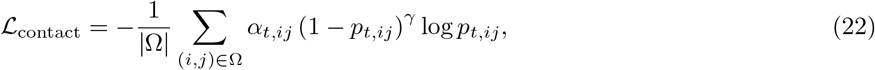

A pointwise projection first reduces the feature dimension, after which a shallow two-dimensional convolutional network incorporates information from neighboring pair positions. This local processing allows the head to recognize spatial patterns such as the short anti-diagonal segments produced by stacked base pairs. A final pointwise projection produces one logit per nucleotide pair, and symmetry is enforced as

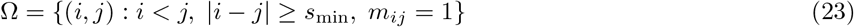

Performance is reported using five complementary measures. *Overall* denotes the F1 score across all predicted and reference base pairs. *PKF* measures recovery of pseudoknot-free base pairs, *CPK* measures recovery of crossing-pseudoknot base pairs, and *PK* measures recovery of base pairs involved in pseudoknots. *Long-distance* measures recovery of reference base pairs satisfying |*i* ™ *j*| ≥ 101. These definitions follow the base-pair-level analysis used by RNA-km.

### C.5 RNA–RNA Interaction Prediction

#### Task definition

MicroRNAs (miRNAs) are small non-coding RNAs of about 22 nucleotides that guide Argonaute proteins to complementary sites in messenger RNAs (mRNAs), most often in the 3^*′*^ untranslated region, leading to translational repression or mRNA decay Bartel (2018). Target recognition depends mainly on pairing between the target site and the miRNA seed region, nucleotides 2–8, and some sites have additional pairing toward the miRNA 3^*′*^ end Bartel (2018). Experimental identification of target sites is costly and time-consuming, so computational prediction is used to prioritize candidate sites for validation Wen et al. (2018). We formulate the task as binary classification: given a miRNA sequence and a candidate target-site sequence, the model predicts whether the two interact. Because the prediction concerns a pair of molecules and depends on their complementarity, this task tests whether representations learned from individual RNAs can support predictions about interactions between RNAs.

#### Dataset

We use the miRNA–target benchmark evaluated by RNAErnie Wang et al. (2024), which RNAErnie refers to as DeepMirTar Wen et al. (2018) and which is distributed as the MirTarRAW dataset. miRNA lengths range from 17 to 26 nucleotides and target-site lengths from 7 to 53 nucleotides. Thymine characters are converted to uracil. Following the RNAErnie implementation, the pairs are shuffled with a fixed random seed, and the first 80% form the training set, which reproduces the RNAErnie training partition. The remaining 20%, which corresponds to the RNAErnie test partition, is divided equally into validation and test sets.

#### Task-specific prediction-head training

The two RNAs are encoded independently using the shared pretrained backbone. Their geometry maps are normalized using the pretraining channel statistics, padded to multiples of the patch size, and processed together with per-nucleotide structural descriptors and validity masks. The backbone’s nucleotide-level readout produces representations 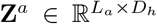 and 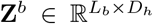, where *L*_*a*_ and *L*_*b*_ denote the effective lengths of the miRNA and target site, respectively. These structural representations are supplied to the prediction head together with one-hot-encoded sequences, nucleotide masks, and the engineered complementarity descriptor.

The default joint-sequence head combines learned base embeddings with the nucleotide-level structural representations. It concatenates the two RNA token streams around a learned separator and processes the resulting joint sequence using a one-dimensional convolution with kernel size 12, followed by a bidirectional LSTM. Masked pooling yields a pair-level representation, which is combined with the engineered complementarity features and passed to a classification network to predict the interaction probability.

An alternative interaction-map head explicitly represents relationships between nucleotide pairs across the two RNAs. It constructs an intermolecular interaction tensor from their nucleotide representations and includes channels indicating Watson–Crick and G·U complementarity. A two-dimensional convolutional network processes this tensor, followed by masked pooling and classification together with the engineered complementarity features. For this head, the backbone readout includes diagonal information, giving *D*_*h*_ = 2*D*; the joint-sequence head uses *D*_*h*_ = *D*. Results are reported in Table S6.

**Table S6:** RNA–RNA interaction prediction performance on the DeepMirTar dataset.

| Method | Accuracy | Recall | Precision | F1 | AUC |
|---|---|---|---|---|---|
| Miranda | 0.659 | 0.652 | 0.666 | 0.659 | 0.687 |
| RNAhybrid | 0.699 | 0.645 | 0.754 | 0.695 | 0.759 |
| PITA | 0.498 | 0.587 | 0.408 | 0.482 | – |
| TargetScan v7.0 | 0.580 | 0.602 | 0.592 | 0.597 | 0.673 |
| TarPmiR | 0.745 | 0.737 | 0.766 | 0.751 | 0.802 |
| DeepMirTar | 0.935 | 0.924 | 0.948 | 0.925 | 0.979 |
| RNABERT | 0.838 | 0.837 | 0.838 | 0.837 | 0.916 |
| RNA-MSM | 0.821 | 0.820 | 0.821 | 0.820 | 0.905 |
| RNA-FM | 0.921 | 0.921 | 0.921 | 0.921 | 0.974 |
| RNAErnie | **0.957** | 0.958 | **0.957** | **0.957** | **0.988** |
| **RNAtomy-FB** | 0.890 | 0.940 | 0.810 | 0.876 | 0.915 |
| **RNAtomy** | 0.942 | **0.961** | 0.824 | 0.887 | 0.957 |

### C.6 RNA–Protein Binding Prediction

#### Task definition

RNA-binding proteins (RBPs) attach to specific sites on their target RNAs and, through these contacts, regulate splicing, polyadenylation, localization, stability, and translation. A systematic census of the human genome identified 1,542 genes encoding RBPs Gerstberger et al. (2014). Their binding sites are mapped experimentally by crosslinking and immunoprecipitation followed by sequencing (CLIP-seq) Licatalosi et al. (2008), but each experiment reports the sites of a single protein in a single cell type. Computational prediction is therefore used to extend these maps to other transcripts and conditions. RBP recognition is determined not only by nucleotide sequence motifs but also by the structural state of the surrounding RNA. In particular, nucleotide accessibility and local base-pairing can influence whether a recognition motif is available for protein binding Sun et al. (2021a).

We formulate RBP–RNA binding prediction as a collection of RBP-specific binary-classification tasks. For each RBP, the model predicts whether a 101-nucleotide RNA window contains a binding site. A separate classifier is trained for each RBP.

#### Dataset

We use the 17 HeLa-cell RBP datasets from the PrismNet benchmark, following the evaluation setting used by RNA-FM and ERNIE-RNA Sun et al. (2021a); Chen et al. (2022); Yin et al. (2025). Each sample is a 101-nucleotide RNA window with a CLIP-seq-derived binary binding label and an experimentally measured icSHAPE reactivity profile Licatalosi et al. (2008); Spitale et al. (2015). The dataset sizes range from 1,398 windows for WTAP to 15,002 windows for the largest RBP dataset, comprising 171,912 windows in total. Thymine characters are converted to uracil. For each RBP, 20% of the samples are reserved for testing, and the remaining 80% are divided 9:1 into training and validation sets. A separate binary classifier is trained and evaluated for each RBP.

#### Task-specific prediction-head training

For each RNA window, the pretrained encoder produces nucleotide-level embeddings

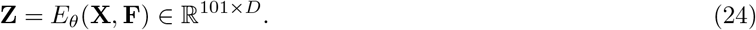

Following the feature-substitution protocol used for the RNA-FM and ERNIE-RNA comparisons, the model embedding replaces the experimental icSHAPE channel in the PrismNet input. The representation supplied to the prediction head at nucleotide *i* is

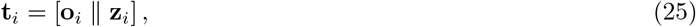

where **o**_*i*_ ∈ ℝ^4^ is the one-hot nucleotide encoding and **z**_*i*_ is the corresponding model embedding. Experimental icSHAPE reactivities are not supplied to the pretrained-model configurations. For **RNAtomy**, the pairwise geometry maps and DSSR descriptors are processed by the pretrained encoder but are not concatenated separately with the input to the prediction head. RealSS+Seq is retained as a reference configuration that combines the sequence with experimentally measured icSHAPE reactivity.

The nucleotide representations are normalized, projected to a common hidden dimension, and processed by a matched PrismNet-based classifier containing convolutional residual blocks and squeeze-and-excitation channel gating. The resulting features are pooled and passed to a multilayer perceptron that produces the binding logit. The same prediction-head design is used for RNA-FM, ERNIE-RNA, and **RNAtomy**, with only the input projection adjusted to accommodate their different embedding dimensions. The results are summarized in Table S7

### C.7 5^′^UTR Mean Ribosome Load Prediction

#### Task definition

Mean ribosome load prediction estimates the average number of ribosomes associated with an mRNA transcript. This quantity reflects translational activity and provides a measure of how efficiently an mRNA is translated. The 5^*′*^ untranslated region can influence translation through its sequence composition, secondary structure, upstream regulatory elements, and interactions with translation-associated factors. We formulate the task as regression, where the model predicts the measured mean ribosome-load value from the mRNA 5^*′*^UTR sequence.

#### Dataset

We use the variable-length 5^*′*^UTR libraries of Sample et al. Sample et al. (2019), in which mean ribosome load was measured by polysome profiling for randomized and human 5^*′*^UTRs of 25 to 100 nucleotides. Following Sample et al., each test set consists of the 100 sequences with the highest read coverage at each length, giving 7,600 random 5^*′*^UTRs (Random7600) and 7,600 human 5^*′*^UTRs (Human7600). The remaining 76,319 random 5^*′*^UTRs are used for model development. Random7600 therefore measures performance on held-out sequences from the training distribution, whereas Human7600 measures transfer to natural human 5^*′*^UTRs.

**Table S7:** RBP–RNA binding prediction performance on the 17 HeLa-cell datasets, measured by AUPRC. A separate classifier is trained for each RBP.

| RBP | Seq<br>(PrismNet) | RealSS+Seq<br>(PrismNet) | RNA-FM+Seq<br>(PrismNet) | ERNIE-RNA+Seq<br>(PrismNet) | **RNAtomy**<br>(PrismNet) |
|---|---|---|---|---|---|
| ELAVL1 | 0.946 | 0.938 | **0.950** | 0.948 | 0.931 |
| UPF1 | 0.933 | **0.956** | 0.934 | 0.943 | 0.949 |
| TIA1 | 0.917 | 0.925 | 0.929 | 0.943 | **0.949** |
| METTL14 | 0.889 | 0.876 | 0.866 | **0.912** | 0.827 |
| HNRNPC | 0.874 | 0.873 | 0.883 | 0.895 | **0.944** |
| CSTF2 | 0.860 | 0.870 | 0.867 | 0.891 | **0.911** |
| TIAL1 | 0.853 | 0.869 | 0.852 | 0.869 | **0.903** |
| U2AF65 | 0.852 | 0.886 | 0.899 | 0.921 | **0.943** |
| EIF4AIII | 0.818 | 0.829 | 0.814 | 0.853 | **0.854** |
| WTAP | 0.804 | 0.843 | 0.801 | **0.876** | 0.725 |
| HNRNPU | 0.798 | 0.821 | 0.829 | 0.849 | **0.864** |
| PTBP1 | 0.793 | 0.840 | 0.799 | 0.839 | **0.903** |
| PTBP1PTBP2 | 0.743 | 0.775 | 0.749 | 0.776 | **0.797** |
| EIF4A3 | 0.738 | 0.758 | 0.751 | **0.834** | **0.834** |
| U2AF2 | 0.712 | 0.764 | 0.705 | **0.807** | 0.701 |
| METTL3 | 0.709 | 0.704 | 0.750 | **0.832** | 0.801 |
| YTHDF2 | 0.610 | 0.629 | 0.638 | **0.725** | 0.700 |

#### Task-specific adaptation and prediction head

For an RNA of length *L*, the pretrained encoder produces a patch-token grid **Z** ∈ ℝ^*G×G×D*^. To obtain a nucleotide-level representation, each patch row is first summarized over its valid columns:

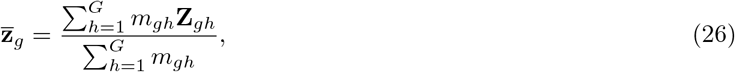

where *m*_*gh*_ indicates whether patch (*g, h*) corresponds to valid RNA positions. The resulting sequence of patch-row representations is then linearly interpolated from *G* patch rows to the original nucleotide length:

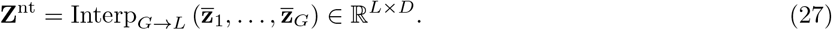

Thus, each nucleotide-level embedding summarizes the interaction profile of its corresponding region across the pairwise map.

For nucleotide *i*, the task-specific input is

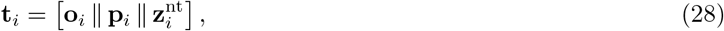

where **o**_*i*_ is the one-hot nucleotide identity and

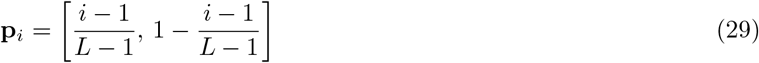

contains normalized positions relative to the cap-proximal and start-codon-proximal ends of the UTR. The pretrained encoder is conditioned on the 23-dimensional DSSR descriptor track through its original conditioning path-way. In an additional ablation, the same DSSR descriptors are concatenated directly with **t**_*i*_ before task-specific processing.

The fused nucleotide tokens are layer-normalized, projected to the task-specific hidden dimension, and processed by four residual one-dimensional convolutional blocks with progressively increasing dilation rates. This design captures both short regulatory motifs and dependencies distributed across the UTR. The resulting tokens are summarized using masked attention, mean, and maximum pooling.

A separate branch encodes six sequence-derived summary variables describing the presence and number of upstream AUG codons, the number aligned with the main coding frame, the normalized position of the first upstream AUG, GC content, and UTR length. Because these variables do not identify a downstream in-frame stop codon, we refer to them as upstream-AUG summaries rather than complete uORF annotations. The pooled nucleotide representation and the upstream-AUG summary embedding are concatenated and mapped to a scalar MRL prediction by a multilayer perceptron.

The primary training objective is mean squared error on standardized MRL:

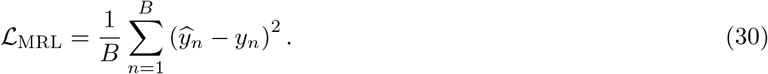

When read-count weighting is enabled, each example is instead assigned a batch-normalized weight proportional to log(1 + *r*_*n*_), where *r*_*n*_ is its total read count:

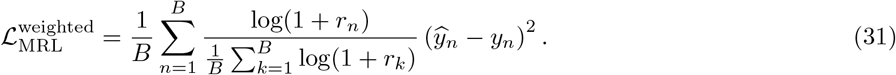

This option gives greater influence to measurements supported by higher sequencing coverage.

### C.8 Internal Ribosome Entry Site Identification

#### Task definition

An internal ribosome entry site (IRES) is an RNA element that recruits the ribosome without the 5^*′*^ cap that most messenger RNAs use to begin translation Weingarten-Gabbay et al. (2016). Circular RNAs have no free ends and therefore no cap, so their translation depends on IRES-like elements, which makes these elements of interest for circular RNA therapeutics Chen et al. (2021); Zhu et al. (2025). Weingarten-Gabbay et al. measured cap-independent translation for tens of thousands of designed sequences drawn from human and viral genomes, and Chen et al. tested this library in circular RNA reporters, finding that structured elements drive much of the translation they observed Weingarten-Gabbay et al. (2016); Chen et al. (2021). We formulate IRES identification as binary classification of a 174-nucleotide sequence as IRES or non-IRES. Because IRES activity depends on RNA structure as well as sequence, this task tests whether structural representations help identify a functional regulatory element.

#### Dataset

We use the IRES benchmark evaluated by structRFM Zhu et al. (2025), which is derived from the circular RNA screen of Chen et al. Chen et al. (2021), with the training and test partition defined by DeepCIP Zhou et al. (2023). We obtained the files from the structRFM data release. The training set contains 12,983 sequences (3,949 IRES and 9,034 non-IRES), and the test set contains 1,164 sequences (582 IRES and 582 non-IRES). All sequences are 174 nucleotides long.

#### Task-specific prediction head

For each 174-nucleotide segment, **RNAtomy** produces a sequence of contex-tualized nucleotide-level embeddings. For nucleotide *i*, the prediction-head input combines its one-hot sequence representation with its **RNAtomy** embedding:

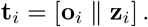

The fused representations are normalized, projected to a shared hidden dimension, and processed using a position-aware residual one-dimensional convolutional network. This architecture retains localized sequence and structural patterns that could be weakened by immediate global pooling. The processed features are summarized using attention and maximum pooling, and a multilayer perceptron maps the resulting RNA-level representation to an IRES classification logit. Raw DSSR descriptors are not concatenated directly to the task-specific head.

We evaluate both frozen-backbone (**RNAtomy-FB**) and fully fine-tuned transfer (**RNAtomy**). The training partition contains substantially more non-IRES than IRES sequences. Following the general balancing strategy used by structRFM Zhu et al. (2025), we train four models using complementary resampled training sets. One member is trained after repeating positive examples three times. Three additional members are trained on independently undersampled datasets, each containing all positive examples and a different equally sized subset of negative examples. Prediction probabilities from the four members are averaged to obtain the final ensemble score.

Each member is optimized using binary cross-entropy:

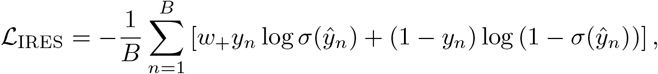

where *ŷ*_*n*_ is the predicted logit and *w*_+_ is calculated from the positive and negative class frequencies of the corresponding resampled training set.

